# Time-based shifts in xylem vulnerability curves of angiosperms based on the flow-centrifuge method

**DOI:** 10.1101/2024.04.02.587697

**Authors:** Luciano M. Silva, Jonas Pfaff, Luciano Pereira, Marcela T. Miranda, Steven Jansen

**Affiliations:** Institute of Botany, Ulm University, Albert-Einstein-Allee 11, 89081 Ulm, Germany

**Keywords:** angiosperm xylem, embolism, hydraulic conductivity, spinning time, water potential, gas diffusion

## Abstract

Centrifuges provide a fast and standard approach to quantify embolism resistance of xylem in vulnerability curves (VCs). Traditionally, embolism formation in centrifuge experiments is assumingly driven by centrifuge speed, and thus pressure, but unaffected by spin time. Here, we explore to what extent embolism resistance is not only pressure but also spin time dependent, and hypothesise that time-stable hydraulic conductivity (K_h_) values could shift VCs. We quantified time-based shifts in flow-centrifuge VCs and their parameter estimations for six angiosperm species by measuring K_h_ at regular intervals over 15 minutes of spinning at a particular speed before a higher speed was applied to the same sample. We compared various VCs per sample based on cumulative spin time, and modelled the relationship between K_h_, xylem water potential (Ψ), and spin time. Time-based changes of K_h_ showed considerable increases and decreases at low and high centrifuge speeds, respectively, which generally shifted VCs towards more positive Ψ values. Values corresponding to 50% loss of hydraulic conductivity (*P*_50_) increased up to 0.72 MPa in *Acer pseudoplatanus*, and on average by 8.5% for all six species compared to VCs that did not consider spin time. By employing an asymptotic exponential model, we estimated time-stable K_h_, which improved the statistical significance of VCs in 5 of the 6 species studied. This model also revealed the instability of VCs at short spin times, and showed that embolism formation in flow-centrifuges followed a saturating exponential growth curve. Although pressure remains the major determinant of embolism formation, spin time should be considered in flow-centrifuge VCs to avoid overestimation of embolism resistance. This spin-time artefact is species-specific, and likely based on relatively slow gas diffusion associated with embolism spreading. It can be minimized by determining time-stable K_h_ values for each centrifuge speed, without considerably extending the experimental time to construct VCs.

## Introduction

Water is mainly transported in vascular plants through the xylem tissue, which provides a system of hollow, non-living conduits, including unicellular tracheids and multicellular vessels (Scholz et al. 2013). During transpiration, the xylem operates under negative pressure, commonly achieving pressure potential values of xylem sap between -1.0 and -3.0 MPa (Tyree et al. 1986). Under conditions of severe drought stress, the physically metastable state of xylem sap in conduits can be disrupted by large gas bubbles, leading to embolism (Tyree et al. 1986, Tyree and Zimmermann 2002).

Embolism resistance of xylem tissue represents a useful trait to understand plant water relations, and has been linked with xylem anatomy, plant physiology (photosynthetic productivity, growth and reproduction), ecological distribution, and drought-induced mortality (Brodribb et al. 2010, 2021, Anderegg et al. 2012, 2016, Choat et al. 2012). Therefore, embolism resistance of conduits holds considerable biological relevance as it determines the hydraulic conductivity of functional xylem, while the loss of a critical amount of conductivity can initiate runaway embolism, resulting in partial or even complete failure of the water transport system (Tyree and Sperry 1988, 1989, Tyree et al. 1998, Hacke and Sperry 2001, Choat et al. 2012). Considering that embolism resistance is a key trait in distinguishing the ability of a species to tolerate drought and frost, it is crucial to have appropriate methods available to quantify embolism resistance at the intra- and interspecific level for a wide range of vegetation types (Duursma and Choat 2016, Sergent et al. 2020).

One common method for assessing embolism levels involves measuring the reduction in hydraulic conductivity (K_h_) when a xylem sample is subjected to decreasing xylem water potential (Ψ) (Sperry 1985). This reduction in K_h_ is typically quantified as the percentage loss of conductivity (PLC), and is expressed with respect to the maximum hydraulic conductivity (K_max_) as follows:

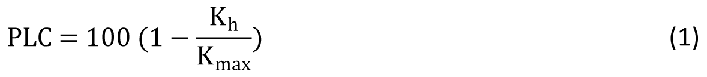

PLC is commonly plotted as a function of Ψ in a vulnerability curve (VC) (Tyree and Sperry 1989). One parameter that is often extracted from these VCs due to their physiological significance in characterizing a species’ embolism resistance is the water potential inducing a 50% loss of conductivity (*P*_50_, MPa), typically located at the steepest part of the VC (Sperry et al. 1988b, Sparks and Black 1999, Domec and Gartner 2001, Sergent et al. 2020). This parameter is generally determined by fitting a response curve (usually sigmoidal, Weilbull or polynomial) to the vulnerability curve data obtained (Cochard 1992, Neufeld et al. 1992, Pammenter and Van der Willigen 1998).

A flow-centrifuge is one of the methods used to construct VCs within a relatively short time frame, typically ranging from approximately 30 minutes to one hour (Cochard 2002, Cochard et al. 2005, 2007). Contrary to the static centrifuge method (Pockman et al. 1995, Alder et al. 1997), the spinning and exposure to negative pressure by the centrifugal forces of a flow-centrifuge are combined with simultaneous measurements of K_h_ (Cochard et al. 2000, Cai and Tyree 2010). The typical procedure for flow-centrifuge VCs involves setting a reference xylem pressure (determined by the rotational speed), usually around −0.1 MPa, to measure the maximum hydraulic conductivity (K_max_). Subsequently, the rotational speed is increased to expose the sample to a more negative pressure. Hydraulic conductivity is measured at least once for each selected pressure level, and the percentage loss of conductivity (PLC) at that pressure is computed using Equation 1. This process is repeated for several pressure levels until the loss of conductivity reaches at least 80% (Sergent et al. 2020).

Centrifuge methods have been used in many studies (Cochard et al. 2013), and assume that embolism formation is driven by the pressure applied, with little or no changes of K_h_ affected by the spin time. Alder et al. (1997) found no significant difference in K_h_ measurements on *Betula occidentalis* when varying the spin time from 3 to 60 minutes. It is generally assumed that embolism formation takes only seconds, suggesting that one minute of sample exposure to a given negative pressure is adequate for precise K_h_ measurements (Cochard et al. 2005). Sergent et al. (2020) proposed exposing a centrifuge sample to a particular speed for three minutes, then taking three K_h_ measurements, and averaging these for an average PLC computation. However, there is accumulating evidence that embolism spreading is at least to some extent not exclusively driven by pressure (Choat et al. 2010, Torres-Ruiz et al. 2015), but also determined by the connectivity and proximity to atmospheric gas (Guan et al. 2021, Avila et al. 2022a), and the timing for embolism to develop (Silva et al. 2023). It has also been shown that K_h_ measurements of cut stem segments may take up to 300 minutes when applying a positive pressure, and that the time required to achieve hydraulic steady state increases with declining xylem water potentials (De Baerdemaeker et al. 2019).

Silva et al. (2023) reported that a sample that was spun for one hour at a fixed Ψ, could exhibit temporal increases in K_h_ ranging between +6% and +40% at low centrifugal speed, but K_h_ may decrease between -41% and -61% at high centrifugal speeds. Interestingly, these changes were significantly larger than previously reported reductions of K_h_ by accumulation of contaminants or gas bubbles on pit membranes (Canny et al. 2007, Espino and Schenk 2011, Krieger and Schymanski 2023). Changes in K_h_ based on the spin time were also in line with evidence that embolism gradually builds up in the centre of centrifuge samples, where the pressure is most negative due to centrifugal forces (Silva et al. submitted). Consequently, considering the spin period component may improve the precision of VCs and associated parameters. Accurate estimations of K_h_ and K_max_ are important to estimate embolism resistance, as K_max_ values measured at low centrifugal speed may increase over time, while values of K_h_ measured at high centrifugal speed may decrease over time (Silva et al. 2023). Decreasing values of K_h_ and/or increasing measurements of K_max_ over time (Equation 1), may significantly change the K_h_/K_max_ ratio. If this would be correct, then such temporal changes may lead to time-based changes of PLC values for a given xylem water potential. If decreasing K_h_ values would become pronounced at high spin speeds, a shift in vulnerability curves (VCs) towards higher Ψ values can be expected, and thus reduced embolism resistance. Except for Alder et al. (1997), which included few measurements only, the relationship between K_h_, Ψ, and spin time remains largely unexplored in centrifuge-based VCs. Wang et al. (2014b) tested the effect of a long (nearly 4 hours), slow spin with a flow-centrifuge on *Robinia pseudoaccacia*, suggesting that increased levels of embolism over time were caused by impurities and/or micro air bubbles as nuclei for embolism. Pit membranes, however, are known not to enable passage of such nuclei or contamination (Zhang et al. 2020, 2024), while dynamic changes in the amount of embolism along centrifuged stem segments cannot be associated with the much faster open-vessel artefact (López et al. 2018, Silva et al. submitted).

Our objective of this paper was to quantify temporal shifts in flow-centrifuge VCs of angiosperms and their parameter estimations resulting from time-dependent phenomena that alter K_h_. This goal was achieved by considering the spin time in VC analyses. We also aimed to model time- stable K_h_ measurements to enhance the reliability of VCs for the flow-centrifuge method. We hypothesised that longer spin times would shift VCs towards higher Ψ values, leading to reduced values of embolism resistance. Furthermore, we expected that these potential VC shifts (including *P*_50_ values) would be species specific, as the variability in K_h_ over time differs among angiosperm species (Silva et al. 2023).

## Material and methods

### Plant material

Samples from six temperate angiosperm tree species (*Acer pseudoplatanus*, *Betula pendula*, *Carpinus betulus*, *Corylus avellana*, *Fagus sylvatica* and *Prunus avium*) were collected between April and June 2023. These are common trees near Ulm University (Germany, 48°25ʹ20.3ʹʹ N, 9°57ʹ20.2ʹʹ E). To avoid an open-vessel artefact (Martin-StPaul et al. 2014), we selected species for flow-centrifuge measurements with average vessel lengths (*L*_V_) shorter than the flow-centrifuge rotor. Vessel length data were based on anatomical data presented in Guan et al. (2022), and based on samples from the same population of tree species at Ulm (Table S1).

For Experiment 1 (temporal dynamics of K_h_), we collected two samples per species for *A. pseudoplatanus*, *B. pendula*, *C. betulus*. For Experiment 2 (spinning time effect on VCs and parameter estimations), we collected four samples (n = 4) per species for *A. pseudoplatanus*, *B. pendula*, *C. betulus*, *C. avellana*, and *F. sylvatica*, and three samples (n = 3) from *P. avium*.

### Sample preparation

Samples were collected around 9:00 AM and were at least 80 cm long. This length significantly exceeded the maximum vessel length of the species studied (Table S1). The branch samples were initially cut in air and promptly submerged in a water-filled bucket. The samples were transported to the lab within 10 minutes, where approximately 10 cm from the stem base was excised underwater. The samples were then immersed in water for over 2 hours to ensure complete rehydration, which prevented embolism formation by cutting branches with xylem sap under negative pressure (Wheeler et al. 2013, Torres-Ruiz et al. 2015, Guan et al. 2021).

After sample rehydration, we selected straight plant segments, with a base diameter (excluding the bark) of 6 ± 2 mm. The samples were recut multiple times underwater to achieve a final length of 27.4 cm (Torres-Ruiz et al. 2015). Then, the ends of the stem segments were trimmed, and the bark was meticulously removed using a sharp razor blade. The prepared stem segments were then mounted in the centrifuge rotor, positioning their ends in cuvettes, while following the natural orientation of xylem sap flow (i.e., with the proximal part of the sample in the upstream reservoir and the distal part in the downstream reservoir). Flow-centrifuge measurements were conducted using a solution of 10 mM KCl and 1 mM CaCl_2_ (Burlett et al. 2022), which was introduced into the cuvettes via a peristaltic pump (Model PP 2201, VWR International bvba, Leuven, Belgium) to measure K_h_.

### Measurements of hydraulic conductivity

Flow-centrifuge measurements were performed with a ChinaTron centrifuge (Model H2100R, Cence Company, Xiangyi, China) with a highly precise control of the rotational speed and temperature (Wang et al. 2014a). Software designed by Wang et al. (2014b) recorded temperature and rotational speed values over time, and calculated the minimum xylem water potential in the middle of a stem segment (Ψ) based on Holbrook et al. (1995). By applying a high-speed camera (scA640-120gm, Basler, Ahrensburg, Germany), the software captured each second potential changes in the position of the gas-liquid meniscus that could be observed inside a cuvette at the upstream stem end. These dynamic changes in the position of the meniscus enabled a precise estimation of K_h_ over time, based on Alder et al. (1997).

Measurements of hydraulic conductivity were started as soon as a desired rotational speed had been achieved. The acceleration and deceleration of the flow-centrifuge were set to 38.5 RPM s^-1^, which provided a useful compromise between reducing the overall spinning time as much as possible, while reducing any potential heating of the sample that would be caused by fast acceleration. Consequently, high spinning speeds required a somewhat longer acceleration time.

*Experiment 1: Investigating temporal dynamics of hydraulic conductivity during one hour of spinning* This experiment aimed to investigate the temporal variation in K_h_ during 1 hour-long spin times. We specifically wanted to test whether the temporal dynamics of hydraulic conductivity for stem samples of *Acer pseudoplatanus*, *Betula pendula*, and *Carpinus betulus* would align with those previously reported for *Corylus avellana*, *Fagus sylvatica*, and *Prunus avium* by Silva et al. (2023). Samples were subjected to a controlled rotational speed and temperature of 22°C while measuring K_h_ over a one-hour duration. Two samples per species were studied. A consistent and low rotational speed of 2000 RPM (equivalent to -0.35 MPa) was applied to all samples, as this speed was considered unlikely to induce embolism. For higher rotational speeds, values between *P*_12_ and *P*_50_ obtained from trial VCs (data not shown) were selected to induce a significant but not fully obstructive level of embolism. Consequently, rotational speeds of 5800, 4000, and 5000 RPM, corresponding to Ψ values of -2.97, -1.41, and -2.21 MPa, were applied to *A. pseudoplatanus*, *B. pendula*, and *C. betulus*, respectively.

K_h_ was assessed multiple times over a duration of one hour. The time interval between consecutive measurements was progressively increased: every 20 seconds during the initial 15 minutes of centrifugation, every minute during the subsequent 15 minutes, and every five minutes for the remaining 15 minutes. A final K_h_ measurement was recorded when the spin time reached the 60- minute mark. This experimental framework was based on Silva et al. (2023), who showed that the most significant changes in K_h_ occurred during a 15-minute time frame after reaching a given rotational speed.

To capture the temporal patterns within individual samples, we employed a time series analysis approach. Considering that absolute values of K_h_ varied across different samples and species, we utilized relative values in addition to absolute values of K_h_. Here, relative change in hydraulic conductivity over time (ΔK_h_, %) was calculated as follows:

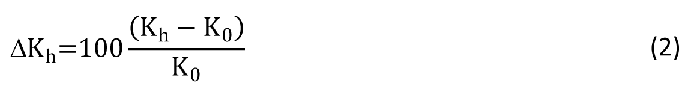

where K_0_ was the hydraulic conductivity measured at time zero, and K_h_ was the current hydraulic conductivity value.

### Experiment 2: Investigating the influence of time on vulnerability curves and parameter estimations

To explore the dynamics between xylem hydraulic conductivity (K_h_), time, and xylem water potential (Ψ) in VCs, we assessed temporal changes of K_h_ at a fixed Ψ before a more negative Ψ was applied. We referred to each of these assessments as a Ψ step. Initially, Ψ for all species was set to - 0.35 MPa, and the initial xylem hydraulic conductivity was measured at regular intervals over a period of 15 minutes. Subsequently, Ψ was decreased to more negative values and maintained for approximately 15 minutes, while K_h_ was measured at regular intervals. The percentage loss of xylem conductivity (PLC) was then determined by equation 1, which considered both the maximum (K_max_) and the actual value of hydraulic conductivity (K_h_). Four to six different Ψ values were applied until a PLC of 90% was reached. The Ψ applied, particularly the more negative ones, varied across the species and were chosen based on previously obtained VCs (data not shown).

For *A. pseudoplatanus*, *B. pendula*, and *C. betulus*, seven K_h_ measurements were taken at each Ψ, resulting in a time interval of 2.5 minutes between measurements. Conversely, for *C. avellana*, *F. sylvatica*, and *P. avium*, four K_h_ measurements were taken at each Ψ, with a 5-minute interval between measurements. Therefore, the dataset resulting from this experiment comprised seven VCs per sample for *A. pseudoplatanus*, *B. pendula*, and *C. betulus*, and four VCs for the species *C. avellana, F. sylvatica*, and *P. avium*. These VCs were constructed by grouping K_h_ measurements based on the duration of spinning. Specifically, one VC was created using data collected immediately after reaching the target Ψ (K_0_). Another VC was generated using K_h_ measurements taken after 5 minutes of spinning, etc. Therefore, for *A. pseudoplatanus*, *B. pendula*, and *C. betulus*, seven VCs were produced with K_h_ measurements taken after 0, 2.5, 5.0, 7.5, 10.0, 12.5, and 15 minutes of spinning at a particular Ψ. Similarly, for *C. avellana*, *F. sylvatica*, and *P. avium*, four VCs were generated for spin times of 0, 5.0, 10.0, and 15 minutes.

Once the number of Ψ applied was between 4 and 6, we presented VCs as a function of the cumulative spin time, which was defined as the total time spent at each pre-set water potential in the flow-centrifuge. Consequently, the time required to reach a target Ψ (i.e., the duration required to increase the rotational speed to achieve a more negative Ψ) was not computed, and this time remained constant across all samples of a particular species. This approach enabled us to analyse how hydraulic conductivity changed over time under varying conditions of Ψ.

A Weibull model was utilized to characterize the relationship between the percentage loss in conductivity (PLC), and Ψ (Neufeld et al. 1992). This model facilitated the estimation of key parameters such as *P*_12_, *P*_50_, and *P*_88_, corresponding to Ψ values at which 12%, 50%, and 88% of conductivity were lost, respectively. To investigate the temporal dynamics of these parameters, we examined their correlation with the cumulative spin time.

Given the variability in *P*_12_, *P*_50_, and *P*_88_ values among different samples and species, both absolute and relative values of these parameters were considered for analysis. Relative values were computed by normalizing *P*_12_, *P*_50_, and *P*_88_ values with their respective initial values (designated as cumulative spin time zero). Relative values were calculated to assess the temporal dynamics of these parameters and to make comparisons across species. For instance, the relative value of *P*_50_ (*P*_50_rel_) was computed as time-based changes in *P*_50_ with respect to the first K_h_ values that were taken immediately after reaching a target Ψ (i.e., K_0_) as reference (i.e., when the cumulative spin time was zero). Therefore, *P*_50___r*el*_ was expressed as:

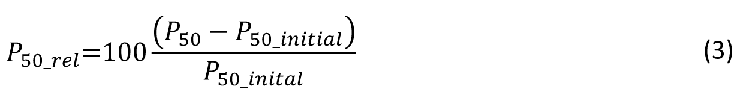

where *P*_50__*_initial_* was *P*_50_ when the cumulative spin time was zero.

The same approach was used to calculated *P*_12*_rel*_ and *P*_88*_rel*_. This normalization procedure allowed for a comparative analysis of the temporal changes in PLC parameters relative to the initial state across different species and experimental conditions.

### The application of an asymptotic exponential model for estimating time-stable hydraulic conductivity

Once K_h_ exhibited significant variation over time and followed a consistent pattern, as described by Silva et al. (2023), it became feasible to characterize this change using an equation model where K_h_ served as the dependent variable and time (t) as the independent variable. The rationale behind this approach was based on the potential to employ a linear or nonlinear regression to calculate the equation coefficients and utilize numerical mathematical solutions to predict time-stable values of hydraulic conductivity. This approach enabled the derivation of more reliable metrics for constructing VCs and reduced the error when making comparisons between different conditions (such as Ψ or temperature), species, or samples.

To estimate a time-stable K_h_, we initially employed an asymptotic exponential model (Goudriaan 1979) in both Experiment 1 and 2 to describe K_h_ as a function of time (K_h_[t]) when Ψ was fixed. The choice of this model was based on Silva et al. (2023) and Miguez et al. (2018), guided by factors such as low error, minimal bias in the distribution of residuals, and the use of the smallest possible number of estimated parameters while meeting regression assumptions. Equation coefficients were determined, and residuals were analysed for trends and outlier removal. In Experiment 2, when no fit was found, we used the average of K_h_ under the same Ψ as time-stable K_h_.

We adopted K_asym_ as the time-stable K_h_ value, which was in the asymptote of the curve, without distinguishing whether K_asym_ corresponded to the lowest or the highest value of hydraulic conductivity. All equations and calculation steps were detailed in the Supporting Information (Methods S1).

### Improving vulnerability curve precision with estimated time-stable K_h_ values

To enhance the reliability of VCs by incorporating stable hydraulic conductivity values over time, we constructed VCs using estimated time-stable K_h_ values derived from the asymptotic exponential model (K_asym_). This methodology was implemented to establish a more robust representation of embolism resistance, in contrast to using only K_h_ measurements immediately taken after reaching a target Ψ (i.e., K_0_). We compared VCs, and evaluated their precision by assessing the R- squared (R^2^) values and computed the Δ*P*_50_, which was the difference between *P*_50_ values calculated using K_asym_ and *P*_50_ values derived from K_0_. Branch samples were incorporated as a random effect. Consequently, all individual data points were combined, and a single fitted curve was plotted for each species using a Weibull model.

### Modelling the relationship between hydraulic conductivity, water potential and time

To integrate the Weibull model that described changes in K_h_ as a function of Ψ with modelled values of K_h_ as a function of time (asymptotic exponential), a comprehensive three-dimensional framework was developed. Initially, for each species, one sample was selected based on the best fit of the asymptotic exponential model, determined by the lowest root mean squared error and satisfying model assumptions. This selection ensured the inclusion of the most representative data for subsequent analyses.

Subsequently, we employed the asymptotic exponential model to simulate changes in K_h_ over a period of 25 minutes for each Ψ step applied to each species. From these simulated K_h_ curves, 400 K_h_ values were extracted at equal intervals of time (3.75 seconds) for each Ψ level. By combining these K_h_ values with corresponding time points across different Ψ levels, 400 VCs were estimated for each species using the Weibull model.

As mentioned above, shifts in *P*_50_ were calculated using the data measured. However, upon integrating the described models, it became feasible to compute *P*_50_ shifts based on model-derived data. This integration yielded a comprehensive 3D curve representing the dynamics of K_h_ over time and Ψ. By extracting *P*_50_ from the 400 modelled vulnerability curves, which compassed the 3D graph (essentially selecting a "slice" of this curve where Ψ equalled the initial *P*_50_), we could illustrate in a curve how *P*_50_ changed over time. This analysis enabled us to investigate whether *P*_50_ increased over time, as hypothesised, and to discern the nature of the relationship, whether it was linear or nonlinear.

### Statistics and data analysis

Data processing, simulations and statistical analyses were performed in RStudio 2022 (version 4.2.2, R Core Team, Boston, USA).

The asymptotic exponential model applied to Experiment 1 and 2 was implemented using the R package, employing the nonlinear least squares method (nls function). Starting values were selected through a grid search or "brute-force" approach using the nls2 package in R. Equation coefficients were determined, and residuals were analysed for trends and outliers. To evaluate the goodness of fit, we used graphical methods and numerical indices as suggested by Miguez et al. (2018). Specifically, when a significant prediction was observed (p-value < 0.05), we calculated the Root Mean Squared Error

(RMSE) and R-squared (R^2^), and visually inspected the standardized residuals to assess the model assumptions of homogeneous variance for the errors.

VCs were fitted to a Weibull function using the R package fitplc (Duursma and Choat 2016), and *P*_12_, *P*_50_ and *P*_88_ values were calculated. To quantify the shifts in absolute and relative values of *P*_12_, *P*_50_, and *P*_88_, we correlated them with the cumulative spin time and considered a significance level of 0.05. This analysis was performed using the lm function within the stats R library.

## Results

### Experiment 1: Investigating temporal dynamics of hydraulic conductivity during one hour of spinning

Both the absolute and relative values of hydraulic conductivity (K_h_ and ΔK_h_) exhibited temporal changes, characterized by increases or decreases depending on the water potential (Ψ) applied (Fig. 1 and Fig. S2). The magnitude and shape of the ΔK_h_(t) curve varied across species. Specifically, both *A. pseudoplatanus* and *B. pendula* demonstrated an exponential curve type, with ΔK_h_ either increasing or decreasing linearly within the first 10 minutes and stabilizing after 20 minutes of spinning (Fig. 1a,b). After one hour of spinning, ΔK_h_ decreased by -39.6% and -38.0% for *A. pseudoplatanus* at Ψ values of -0.4 and -3.0 MPa, respectively, while it increased by 20.6% and decreased by -17.8% for *B. pendula* at Ψ values of -0.4 and -1.4 MPa, respectively (Fig. 1a,b). Conversely, *C. betulus* maintained relatively stable values, decreasing by -0.9% and -4.5% at Ψ values of -0.4 and -2.2 MPa, respectively (Fig. 1c). *C. betulus* curve profile resembled a sum of exponential equations, with ΔK_h_ increasing in the first 5 minutes followed by a subsequent decline to negative values.

**Fig. 1.**
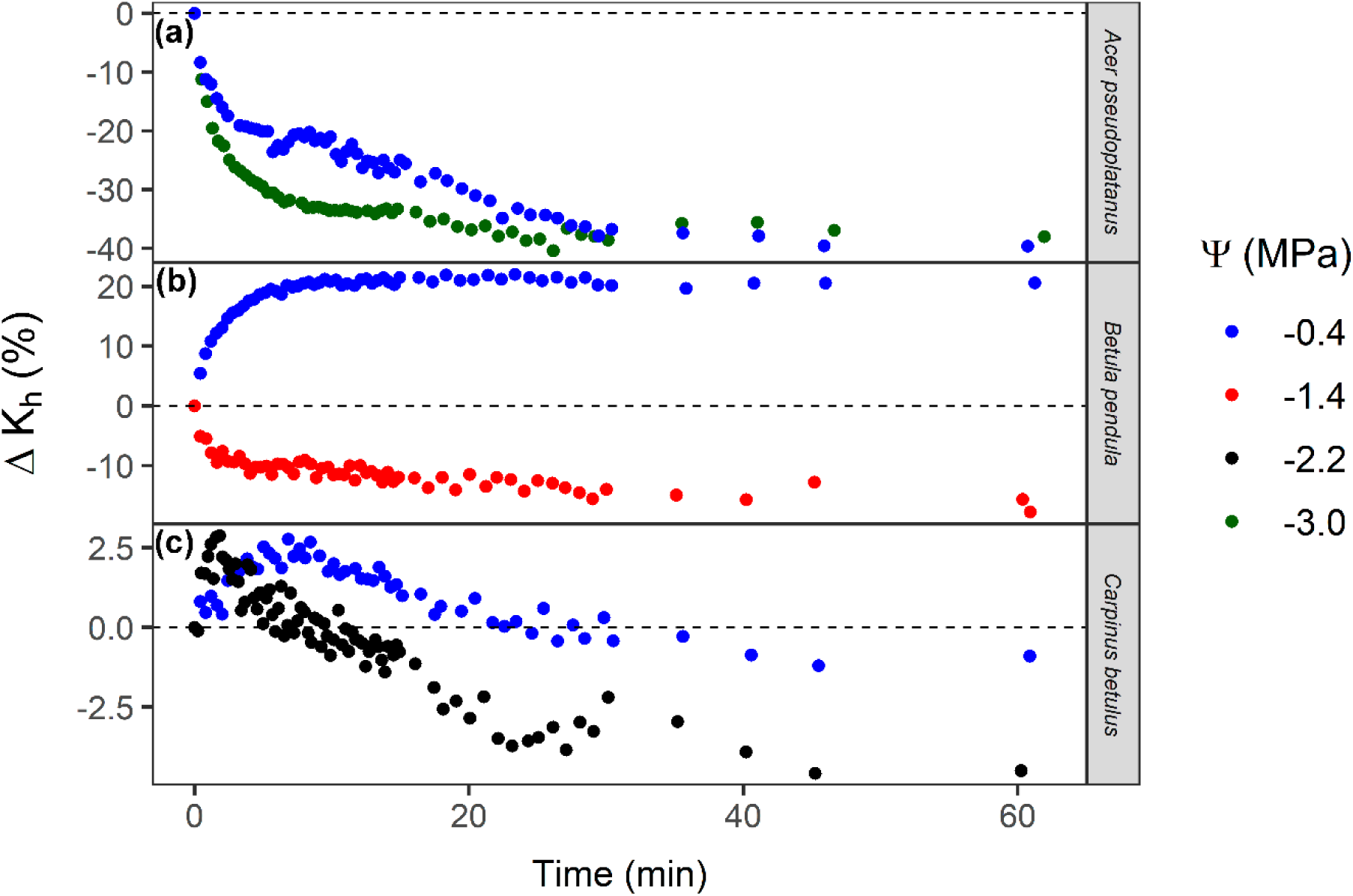
Changes in the hydraulic conductivity over time (ΔK_h_) for stem samples of three angiosperm species (*A. pseudoplatanus*, *B. pendula*, and *C. betulus*), which were spun in a flow-centrifuge for 1 hour and temperature of 22°C. The samples were subjected to a constant xylem water potential (Ψ) based on the rotational speed applied. ΔK_h_ is a relative value calculated through equation 2. Each colour represents a separate sample that was spun at a fixed speed.

The asymptotic exponential model demonstrated a significant prediction of K_h_ values (p-value < 0.05) for *A. pseudoplatanus*, *B. pendula*, and *C. betulus*, covering all Ψ levels (Fig. S2). The model’s effectiveness was evident through low Root Mean Squared Error (RMSE) values and R-squared (R^2^) values close to 1, indicating precise predictions with minimal error. Examination of standardised residuals revealed a random distribution, particularly noticeable for *B. pendula* at Ψ values of -1.4 MPa and C. *betulus* at Ψ values of -0.4 and -2.2 MPa (Fig. S3c,e,f). Although residual patterns exhibited wave-like sinusoidal shapes in other instances, there was no discernible trend of model underestimation or overestimation for low or high yield predictions. Therefore, the asymptotic exponential model was found to be a robust method for characterizing the changes in K_h_ over time.

*Experiment 2: Investigating the influence of time on vulnerability curves and parameter estimations* The percentage loss of xylem conductivity (PLC) was primarily driven by the water potential (Ψ) applied, with PLC increasing as Ψ became more negative (Fig. 2). However, time also played a significant role. Even at a fixed Ψ, PLC exhibited a logarithmic increase or decrease over time, depending on the magnitude of Ψ values. Overall, PLC decreased over time when Ψ was -0.4 MPa. The most pronounced decrease in PLC at a fixed Ψ during the 15-minute K_h_ measurements was observed in *P. avium* (sample 2, Fig. 2v) at a Ψ of -0.4 MPa, where PLC decreased from +30.2% to 0%. Conversely, the most evident increase in PLC was observed in *B. pendula* (sample 4, Fig. 2h) at a Ψ of -1.6 MPa, with PLC increasing from +36.5% to +60.4%. *C. betulus* exhibited the most stable PLC values over time, with a maximum decrease from +7.7% to +0.6% at a Ψ of -0.8 MPa (sample 1, Fig. 2i) and a maximum increase from +50.8% to +63.1% at Ψ of -3.2 MPa (sample 4, Fig. 2l).

**Fig. 2.**
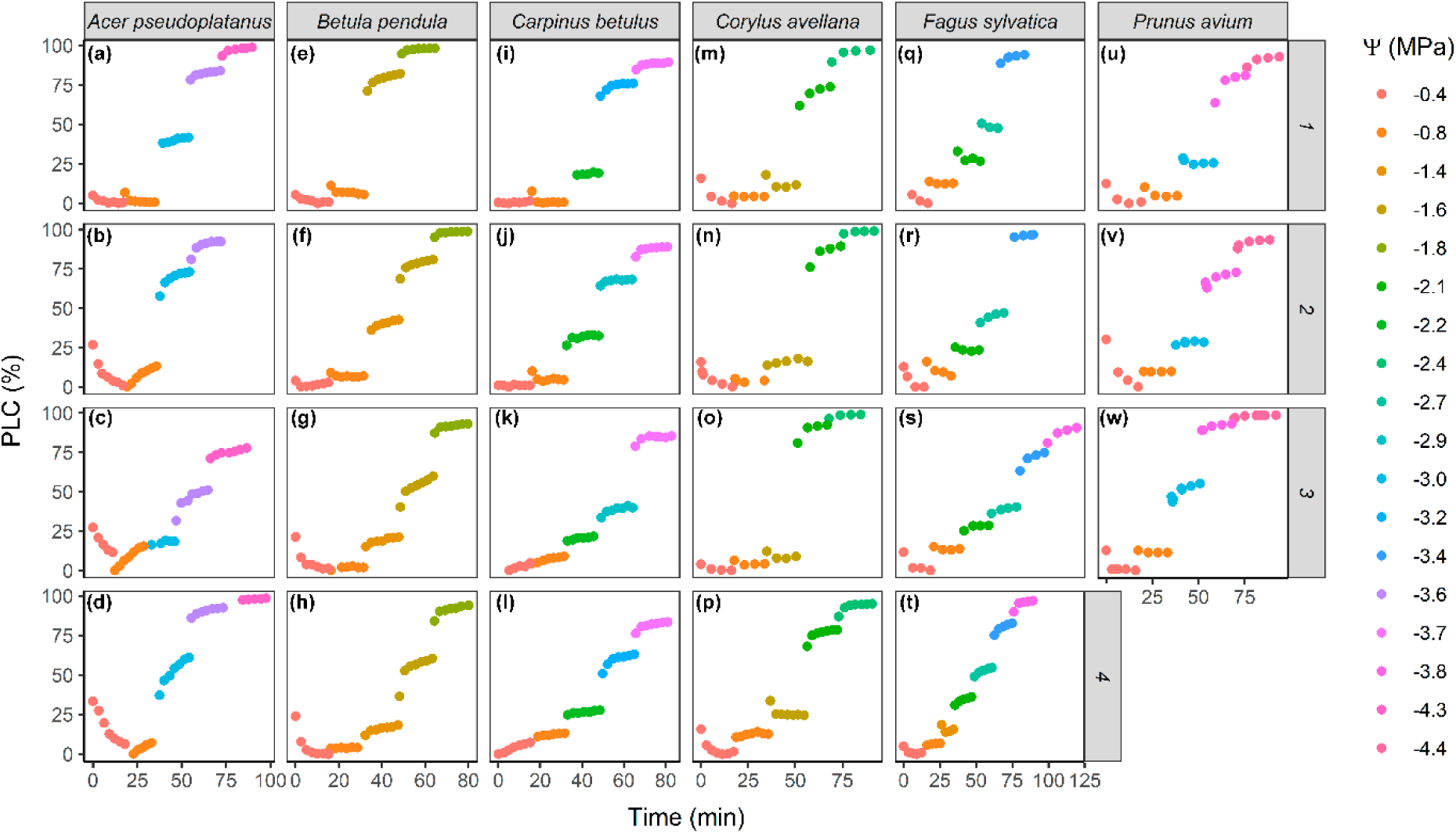
Changes in the percentage loss of xylem hydraulic conductivity (PLC) over time and water potential (Ψ). Stem samples of six angiosperm species were spun in a flow-centrifuge at a constant temperature of 22°C and subjected to different Ψ in the centre of the sample, based on the rotational speed applied. Each colour represents a different Ψ. The figure presents a grid layout with PLC plotted as a function of time (in minutes) for each species (columns) and sample (rows), with n = 4 for *Acer pseudoplatanus*, *Betula pendula*, *Carpinus betulus*, *Corylus avellana*, and *Fagus sylvatica*, and n = 3 for *Prunus avium*.

Overall, when Ψ resulted in PLC of +25% or higher, PLC tended to increase over time. However, when the Ψ applied resulted in PLC lower than 25%, the results varied. For instance, PLC increased over time for *C. betulus* in sample 3 and 4 (Fig. 2k,l), remained stable for the same species in sample 1 and 2 (Fig. 2i,j), and decreased for *P. avium* and *F. sylvatica*. Notably, for *A. pseudoplatanus* (sample 2, 3, and 4; Fig. 2b,c,d), even changing Ψ to a more negative value (from -0.4 to -0.8 MPa) resulted in a decrease in PLC.

The vulnerability curves (VCs) depicted in Fig S4 revealed a significant influence of time even within a single sample, as evidenced by changes in slope and curve shift. Particularly notable for *A. pseudoplatanus* (sample 2, Fig. S4b) and *F. sylvatica* (sample 3, Fig. S4s), a longer cumulative spin time tended to transform the curve into an exponential shape. Overall, increasing cumulative spin time, which represented the total spin time of all Ψ steps, shifted a VC towards a slightly more positive direction of Ψ, leading to an increase in *P*_50_ over time.

We observed a strong and positive correlation between the absolute values of *P*_50_ and cumulative spin time (Fig. 3). This correlation proved statistically significant (p < 0.05) for the majority of the samples and species, with exceptions noted for *A. pseudoplatanus* (sample 3, Fig. 3a), *C. betulus* (samples 3 and 4, Fig. 3c), *F. sylvatica* (sample 1 and 3, Fig. 3e), and *P. avium* (sample 1, Fig. 3f). When significant, the correlation between *P*_50_ and cumulative spin time exhibited high values of R-squared (R^2^), ranging from 0.69 to 0.99.

**Fig. 3.**
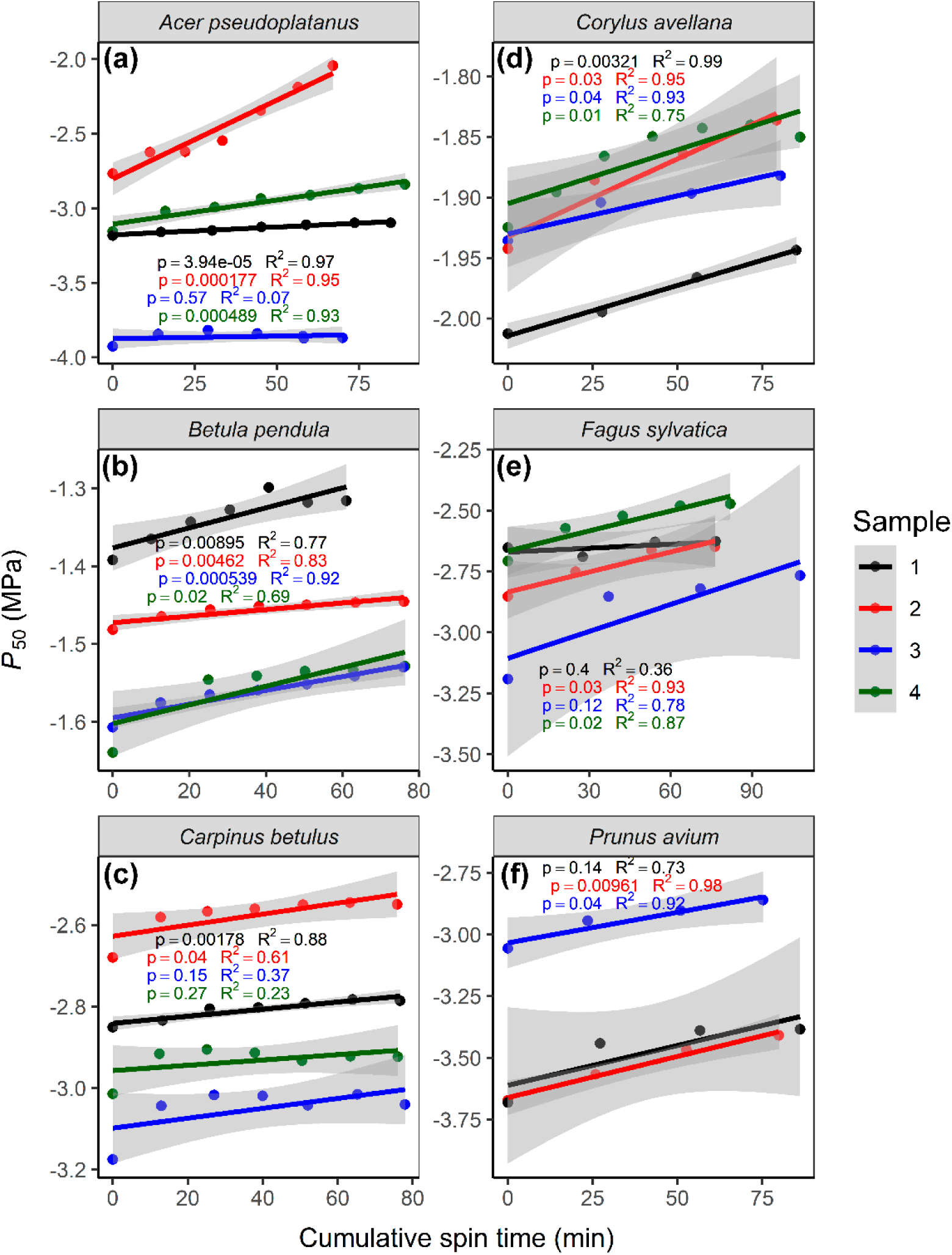
Absolute water potential values corresponding to 50% loss in hydraulic conductivity (*P*_50_) as a function of cumulative spin time for six angiosperm species. Cumulative spin time is calculated as the accumulated time spent at each pre-set centrifuge speed. Each colour represents a different repetition, with n = 4 for *Acer pseudoplatanus*, *Betula pendula*, *Carpinus betulus*, *Corylus avellana*, and *Fagus sylvatica*, and n = 3 for *Prunus avium*. Points represent estimated *P*_50_ values obtained from vulnerability curves in Fig S4. Coloured lines indicate the correlation between *P*_50_ and cumulative spin time, with corresponding confidence intervals in grey, and p-values (p) and R-squared (R^2^) values provided.

When relative values of *P*_50_ were combined across all samples from the same species for comparative analysis, we observed a significant positive correlation between relative values of *P*_50_ and cumulative spin time across all species (Fig. 4). However, the species studied diverged regarding the strength of *P*_50_’s correlation with time and the rate of change in both absolute and relative values of *P*_50_. This correlation was strong for *P. avium* (R^2^ = 0.81; Fig. 4f) and weak for *A. pseudoplatanus* (R^2^ = 0.16; Fig. 4a). Notably, *A. pseudoplatanus* exhibited the most significant increase in *P*_50_, by 0.72 MPa or 26%, in terms of both absolute and relative values, respectively, when the cumulative spin time was 76 minutes. Conversely, *C. betulus* demonstrated the most stable *P*_50_ values over time, increasing by only 0.04 MPa or 4.9% when the cumulative spin time was 67 minutes.

**Fig. 4.**
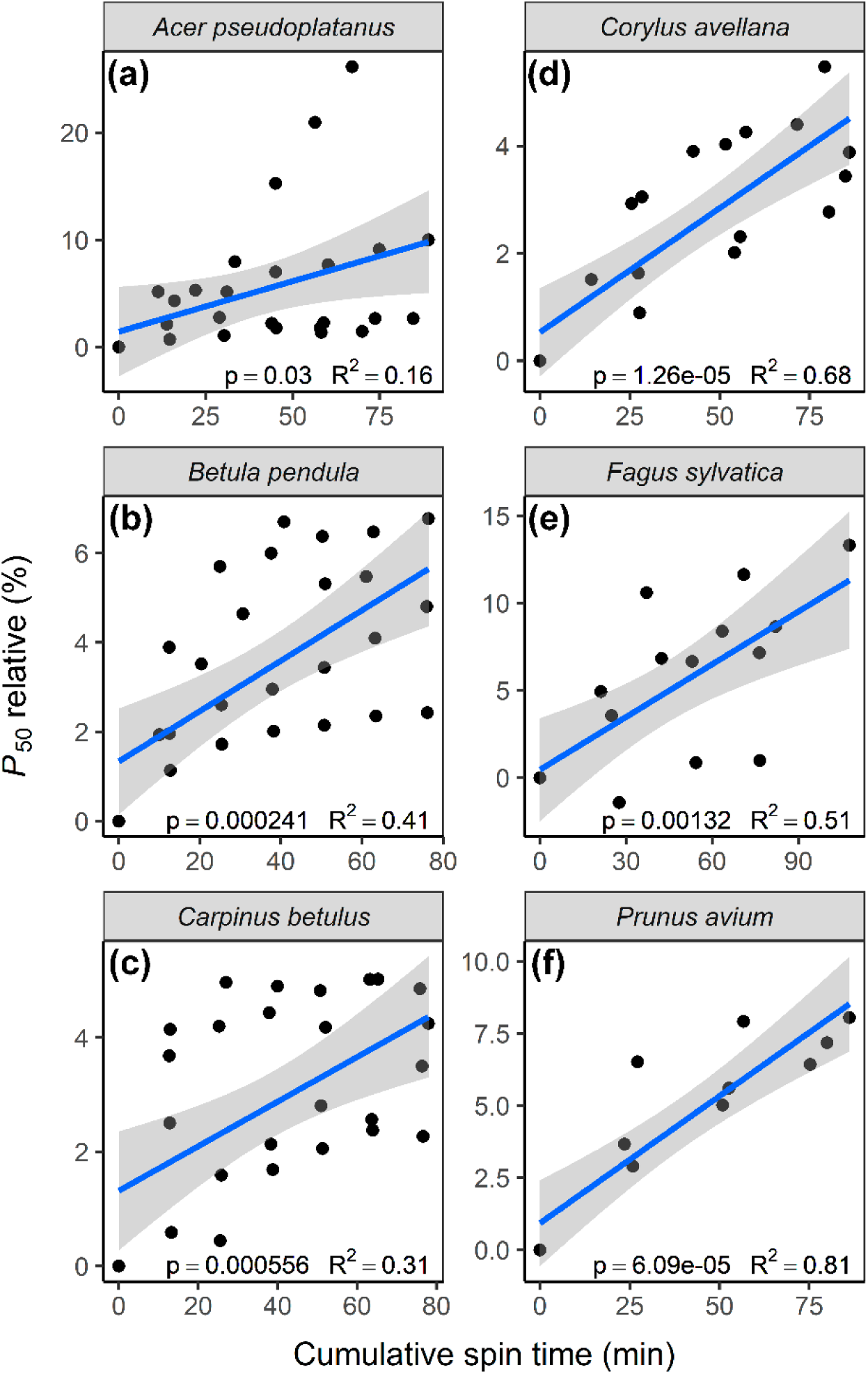
Relative water potential values corresponding to 50% loss in hydraulic conductivity (*P*_50_) as a function of cumulative spin time for six angiosperm species (*Acer pseudoplatanus*, *Betula pendula*, *Carpinus betulus*, *Corylus avellana*, *Fagus sylvatica*, and *Prunus avium*). Cumulative spin time is calculated as the accumulated time spent at each pre-set centrifuge speed. Points represent the relative values of *P*_50_ that were calculated through equation 3 and represent the temporal changes in *P*_50_ relative to its value when cumulative spin time was equal to zero. The blue line indicates the correlation between *P*_50_ relative and cumulative spin time, with corresponding confidence intervals in grey, and p-value (p) and R-squared (R^2^) values provided.

On average, the relative *P*_50_ values increased by 9.8%, 5.6%, 4.2%, 4.5%, 11.3%, and 8.5% for *Acer pseudoplatanus*, *Betula pendula*, *Carpinus betulus*, *Corylus avellana*, *Fagus sylvatica*, and *Prunus avium*, respectively (Fig. 4).

Absolute and relative values of *P*_88_ exhibited a similar behaviour as *P*_50_: absolute values of *P*_88_ showed a significant positive correlation (p < 0.05) with cumulative spin time for most species and samples (Fig S5a), while relative values of *P*_88_ (combined across all samples from the same species) correlated significantly (p < 0.05) for all species (Fig S6a). On average, relative *P*_88_ increased by 6.3%, 7.9%, 9.0%, 8.7%, 8.0%, and 8.0% for *A. pseudoplatanus*, *B. pendula*, *C. betulus*, *C. avellana*, *F. sylvatica*, and *P. avium*, respectively.

In contrast, absolute values of *P*_12_ did not exhibit a significant correlation with cumulative spin time for most samples (Fig S5b); significant and positive correlations were observed only for *A. pseudoplatanus* (samples 2 and 4), *C. avellana* (sample 2), *F. sylvatica* (sample 2 and 4), and *Prunus avium* (sample 2 and 3). Notably, for *C. betulus* (sample 4), *P*_12_ showed a strongly negative and significant correlation with cumulative spin time. Consequently, relative values of *P*_12_ did not display a significant correlation with cumulative spin time across species, and significant, average increases were found for *F. sylvatica* (15.9%) and *P. avium* (9.1%) (Fig S6b).

Combining the relative values of *P*_12_, *P*_50_, and *P*_88_ across all species and samples revealed a positive and significant correlation with cumulative spin time (p < 0.5; Fig. 5). The *P*_12_ values exhibited a weak correlation (R^2^ = 0.05), and increased on average by 6.3% (Fig. 5a). *P*_50_ values showed a stronger correlation (R^2^ = 0.26) and increased on average by 8.5% (Fig. 5b). *P*_88_ demonstrated the strongest correlation among these parameters (R^2^ = 0.34) and increased on average by 9.5% (Fig. 5c).

**Fig. 5.**
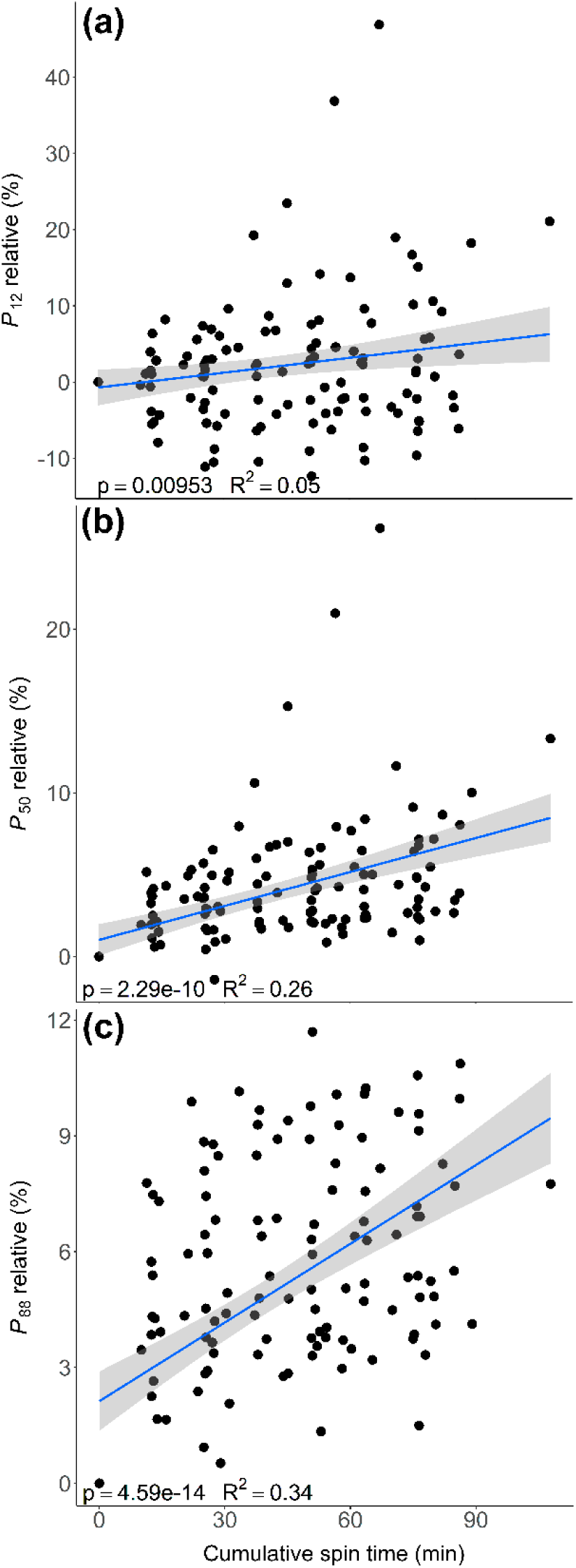
Relative water potential values corresponding to 12%, 50% and 88% loss in hydraulic conductivity (*P*_12_, *P*_50_ and *P*_88_, respectively) as a function of cumulative spin time. Cumulative spin time is calculated as the accumulated time spent at each pre-set centrifuge speed. Points represent the relative values of *P*_12_, *P*_50_ and *P*_88_ that were calculated through equation 3 and represent the temporal changes in these parameters relative to their values when cumulative spin time was equal to zero. The blue line indicates the correlation between relative *P*_50_ values and cumulative spin time, with corresponding confidence intervals in grey, and p-values (p) and R-squared (R^2^) values provided.

### The application of an asymptotic exponential model for estimating time-stable hydraulic conductivity

The asymptotic exponential model was able to predict K_h_ values for the majority of samples across all species and conditions (Fig. S8, S9, S10, S11, S12 and S13). Out of 116 K_h_(t) curves, the model achieved 100 significant predictions (p-value < 0.05).

However, in the remaining 16 instances, there was a lack of fitting due to various reasons. For *F. sylvatica* (sample 1, Fig. S12) and *B. pendula* (sample 2, Fig. S9) at a Ψ of -0.4 MPa, we were unable to fit the model accurately due to the lack of a plateau, indicating that the curve was in the initial phase where K_h_ either increased or decreased linearly over time. For the remaining 14 cases, the lack of an initial slope or the random distribution of K_h_ values prevented us from fitting the model. The limited number of four K_h_ measurements for *C. avellana*, *F. sylvatica*, and *P. avium* was associated with the lack of fitting, as these species collectively contributed to 12 curves without significant fitting Fig. S11, S12 and S13).

### Improving vulnerability curve precision with estimated time-stable K_h_ values

A considerable improvement of the VCs was obtained when we compared VCs based on the estimated time-stable K_h_ values, which were derived from the asymptotic exponential model (K_asym_), against VCs based on K_h_ measurements immediately taken after reaching the target Ψ (K_0_; Fig. 6). K_asym_ curves exhibited higher R^2^ values for 5 out of the 6 species studied, suggesting a closer fit to the Weibull model. The most pronounced improvement was observed for *B. pendula*, where the final plateau at high levels of PLC was more pronounced, leading to an increase in the R^2^ value from 0.90 to 0.97 (Fig. 6b). Overall, employing K_asym_ resulted in higher *P*_50_ values, indicating a shift of the VC to less negative Ψ values. The variation of *P*_50_ (Δ*P*_50_) was maximal for A*. pseudoplatanus* (0.33 MPa; Fig. 6a) and minimal for both *B. pendula* and *C. betulus* (0.06 MPa; Fig. 6c).

**Fig. 6.**
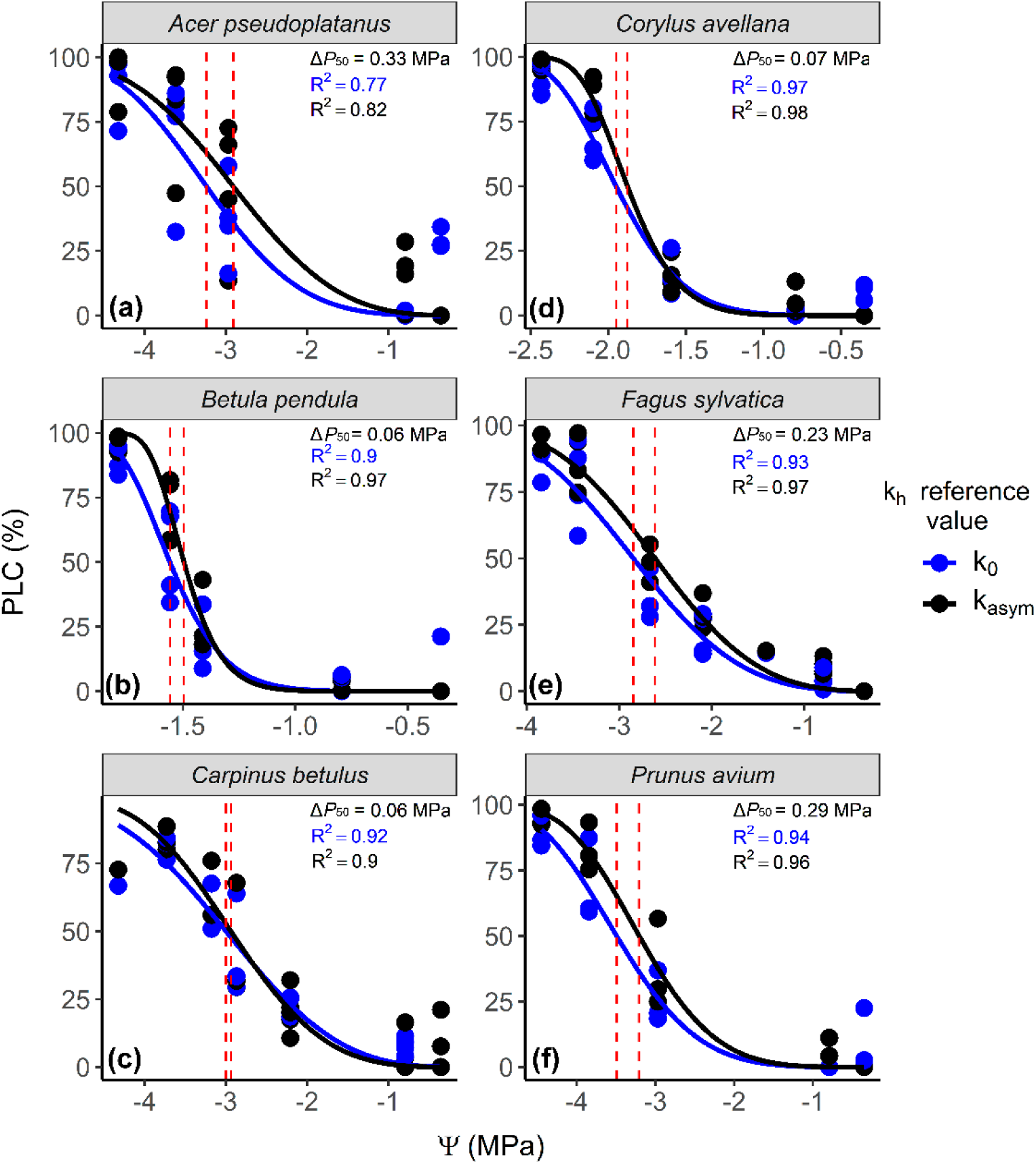
Comparison of vulnerability curves using estimated time-stable hydraulic conductivity values derived from the asymptotic exponential model (K_asym_, black dots) and hydraulic conductivity measurements taken immediately after reaching the target water potential (K_0_, blue dots) for six angiosperm species. Both black and blue dots represent the percentage loss of xylem conductivity (PLC) as function of water potential (Ψ) with all samples of each species combined to generate a single vulnerability curve. The curves represent vulnerability curves predicted by the Weibull model with their respective R-squared (R^2^) values. Dashed vertical red lines indicate the Ψ corresponding to 50% loss in conductivity (*P*_50_). Δ*P*_50_ denotes the difference between *P*_50_ values calculated using K_asym_ and those derived from K_0_.

### Modelling the relationship between hydraulic conductivity, water potential and time

We selected one sample as the most representative data for modelling the relationship between hydraulic conductivity, water potential, and time, namely samples 2, 2, 5, 4, 3 and 1 from *A. pseudoplatanus, B. pendula, C. betulus, C. avellana, F. sylvatica, and P. avium*, respectively (Fig. 7). This integrative model revealed that VCs were unstable at cumulative spin times close to zero. Specifically, the curves tended to shift towards more positive Ψ values for all species, and the slope at *P*_50_ values tended to increase for *C. betulus, C. avellana,* and *P. avium* (Fig. 7c,d,f). After 100 minutes of cumulative spin time, the VCs tended to be less variable, and were more sigmoidal in shape. Particularly for *A. pseudoplatanus*, longer cumulative spin times led to changes in the shape and slope of its VCs, resulting in considerable increases in our *P*_50_ estimation (Fig. 7a).

**Fig. 7.**
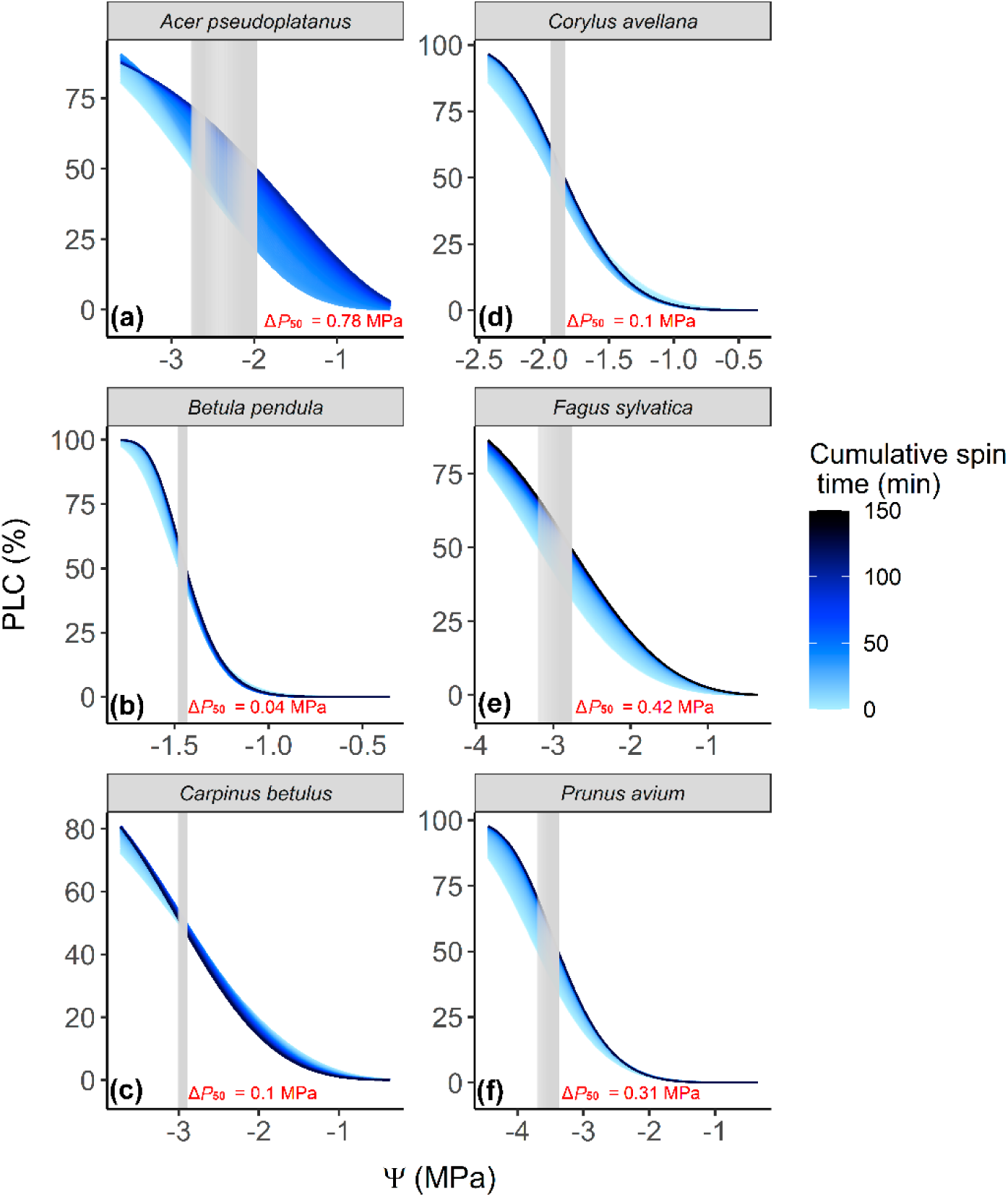
Modelled relationship between the percentage loss of xylem hydraulic conductivity (PLC), water potential (Ψ), and cumulative spin time for six angiosperm species. For each species, 400 vulnerability curves were estimated using the Weibull model after simulating hydraulic conductivity dynamics for 25 minutes at each Ψ step using the asymptotic exponential model. Curves are colour-coded according to the cumulative spin time, which is calculated as the accumulated time spent at each pre-set centrifuge speed in a flow-centrifuge. Vertical grey lines represent a shift in the Ψ corresponding to a PLC of 50%, numerically represented in the plot as Δ*P*_50_.

While we observed an increase in *P*_50_ for 5 out of 6 species, the relationship between *P*_50_ and cumulative spin time was not linear (Fig. 8). *For B. pendula, C. avellana, F. sylvatica,* and *P. avium*, we observed a saturating exponential growth curve type for the relationship between *P*_50_ and cumulative spin time, characterized by initial linear growth followed by temporal stabilization. For *C. avellana*, we observed an initial slight decrease in *P*_50_ from time 0 to 6 minutes before values increased (Fig. 8d). In the case of *A. pseudoplatanus*, the exponential curve was not well defined, with an inflection point at a cumulative spin time of 25 minutes (Fig. 8a). *C. betulus* was the only species that displayed a unique pattern, with *P*_50_ values first increasing for a cumulative spin time between 0 and 34 minutes, and then decreasing values of *P*_50_ (Fig. 8c). Overall, *A. pseudoplatanus, B. pendula, C. betulus, C. avellana, F*.

**Fig.8.**
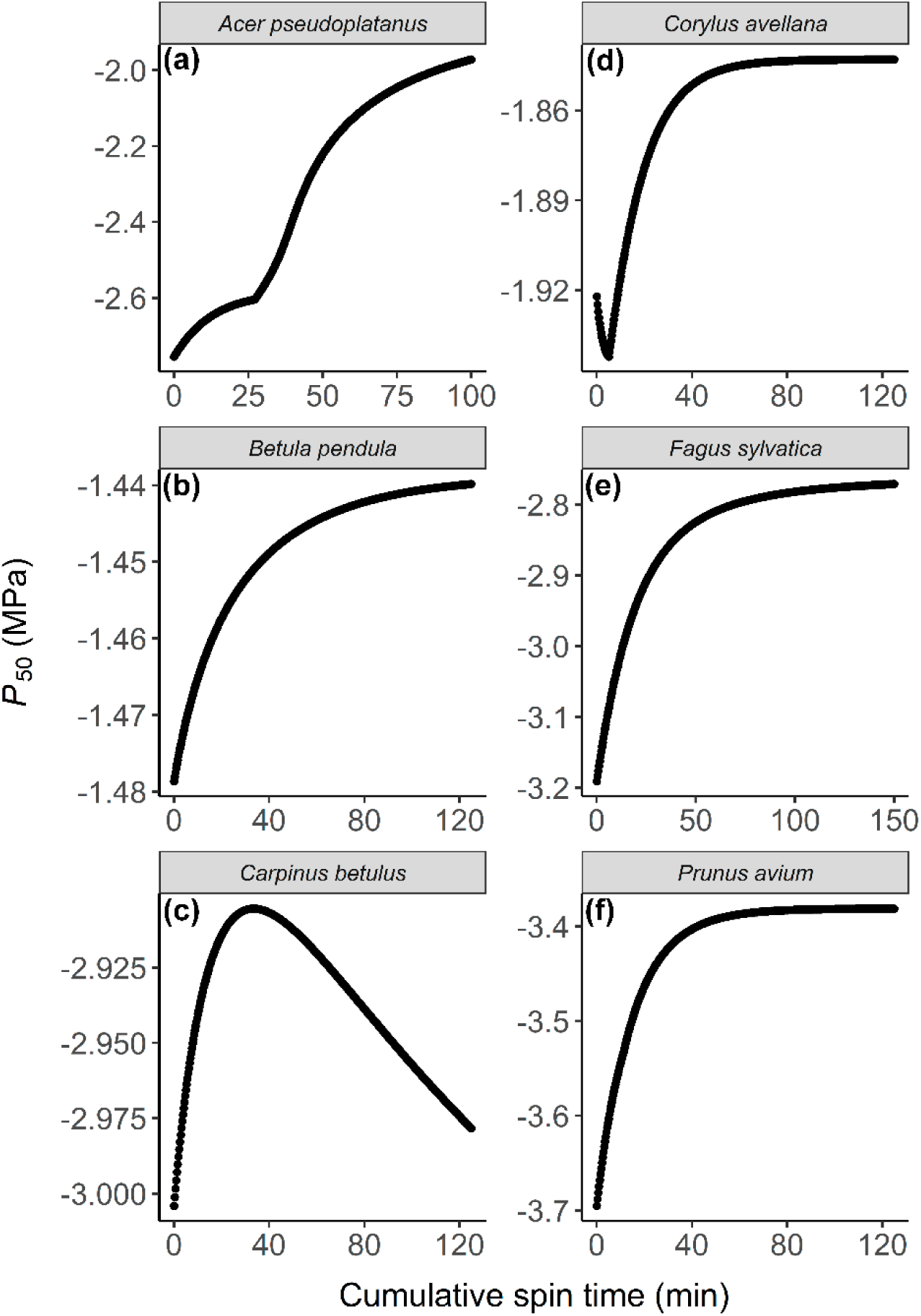
Time-based shifts in water potential values corresponding to 50% loss in hydraulic conductivity (*P*_50_) for six angiosperm species. The points represent *P*_50_ shifts based on data derived from a combination of the Weibull and asymptotic exponential models, obtained from vulnerability curves in Fig. 7. Cumulative spin time is calculated as the accumulated time spent at each pre-set centrifuge speed in a flow-centrifuge.

*sylvatica,* and *P. avium* exhibited modelled increases in *P*_50_ of 0.78, 0.04, 0.10, 0.10, 0.41, and 0.31 MPa, respectively.

## Discussion

By incorporating time into vulnerability curves (VCs), we confirmed our hypothesis that flow-centrifuge VCs shift towards higher positive water potential (Ψ) values over time. Although pressure represents by far the most important determinant, these minor shifts significantly impacted VC parameter estimations, such as the *P*_12_, *P*_50_, and *P*_88_ values (Fig. 3, 4, 5, S5 and S6). Moreover, the modelled data suggest that this time-dependent effect on VCs might not be linear (Fig. 8). These findings reflect earlier suggestions about the dynamic, instable behaviour of hydraulic conductivity (K_h_) over time, as observed in our Experiment 1 (Fig. 1; Silva et al. 2023). Our analyses revealed that time-based measurements increase the gap between K_h_ and maximum hydraulic conductivity (K_max_): values of K_max_ measured at a low centrifugal speed appear to increase over time, whereas K_h_ values measured at a high centrifugal speed tend to decrease.

The increase in K_h_ over time has been attributed to various processes, including the rearrangement and/or dissolution of embolism within xylem conduits (Sperry et al. 1988a, De Baerdemaeker et al. 2019), the ionic effect (Jansen et al. 2011), bubble dissolution (Yang and Tyree 1992), and even osmotic water uptake by cells, or capillary uptake by the apoplast (Taneda and Sperry 2008, Torres-Ruiz et al. 2012). However, this phenomenon has rarely been observed at negative pressure potentials, suggesting that it either seldomly occurs, or that the processes leading to a decrease in K_h_ at more negative Ψ are stronger than those increasing K_h_. Regardless of the specific mechanisms driving temporal increases of K_h_, we argue that this increase in K_h_ should be quantified and taken into account for vulnerability curves (VCs), given its significant influence on estimating maximum hydraulic conductivity (K_max_) at reference xylem pressure. The practical implication of time- stable K_max_ values is that it enables us to obtain more accurate VCs, with a more adequate curve fitting. The decrease in K_h_ over time can be attributed to embolism, in addition to various other phenomena. Generally, these are commonly referred to as a ‘natural’ decline in K_h_, typically associated with pit membrane clogging, wound response, radial or background flow, or compression of intervessel pit membranes (Espino and Schenk 2011, De Baerdemaeker et al. 2019). However, Silva et al. (2023) suggested that this ‘natural’ decline in K_h_ over 8 hours is very slow, limited to ca. 0.7%, and asserted that the much higher decreases in K_h_ over shorter time periods (minutes) are due to gradual embolism formation in a flow centrifuge. Therefore, embolism spreading is likely the driving force behind the high decreases in K_h_ as observed in Experiment 1, which ranged from 4.5% to 39.6%. Evidence for gradual embolism formation was also confirmed by carefully analysing the spatial distribution and temporal changes of embolised vessels along centrifuge samples (Silva et al. submitted).

Although a comprehensive mechanistic understanding of how embolism propagates between vessels remains elusive (Jansen and Schenk 2015, Lens et al. 2022, Lintunen et al. 2022), embolism spreading among conduits has been attributed to gas diffusion (Wang et al. 2015, Yang et al. 2022, Pereira et al. 2023, Silva et al. 2023). Since gas diffusion within the xylem can be relatively slow, ranging from seconds for axial diffusion to many hours for radial diffusion (Sorz and Hietz 2006, Yang et al. 2022), such association between embolism spreading and gas diffusion could elucidate why the time component is important in K_h_ decreases at the high-speed in a flow-centrifuge.

Considering that at least a certain amount of embolism formation requires time to develop, spinning duration may especially influence the slope of VCs, resulting in a steep slope for long spin times, and a low slope for a short spin duration. Nonetheless, no definitive pattern emerged when correlating slope and cumulative spin time (Fig. S7), with few samples showing a significant increase in slope with cumulative spin time. The observed variability of these parameters could be attributed to the influence of K_max_ on the slope. While Yin et al. (2019) asserted that the slope of VCs generated by the flow-centrifuge method is notably steeper than those produced by other methods, we argue that their analysis did not fully consider the VC dynamics presented here.

Assuming refilling (Taneda and Sperry 2008) as the process responsible for increasing K_h_ (Taneda and Sperry 2008), and embolism spreading (Silva et al. 2023) as the process that decreases K_h_ over time (Silva et al. 2023), we can speculate how their interaction shapes the various curve types observed in experiments 1 and 2, as well as in our model, delineating the temporal dynamics of K_h_. Taking *Carpinus betulus* as an example, which presented a distinct curve type in both the measured and modelled data (Fig. 1c and 8c, respectively), we observe an initial exponential increase in K_h_. This increase is most likely driven by vessel refilling, where water influx into the xylem conduits outpaces the rate of embolism spreading. This refilling process drives an accelerating change in K_h_. However, as spin time progresses, embolism appears to become gradually more pronounced than refilling. This transition is demarcated by an inflection point in the curve. Consequently, the curve experiences a distinct reversal, marked by an exponential decline in K_h_ over time, as embolisms spread further, diminishing hydraulic conductivity. This observed pattern signifies a dynamic interplay between vessel refilling and embolism spread, with an initial phase dominated by refilling, and then a phase when embolism spreading prevails and K_h_ values decline. This interpretation would also explain findings reported by Silva et al. (2023).

An interplay between processes influencing K_h_ over time might may explain the results obtained in this study. At Ψ values near zero and at extremely negative values, one phenomenon is likely to have a dominating effect on K_h_, while the impact of other K_h_-related processes become negligible. Specifically, at Ψ near zero, refilling prevails, as the xylem tension at this point is unlikely to induce embolism in most vessels. Conversely, under significantly negative Ψ conditions, pressure-driven embolism spreading becomes most dominant. In this scenario, we can simplify the changes in K_h_ over time using a single exponential equation, such as the asymptotic exponential, which showed to have a significant predictive power for most samples in Experiment 1 and 2 (Fig. S2, S8, S9, S10, S11, S12 and S13). The asymptotic exponential model is often employed to describe biological phenomena, primarily due to its exponential saturating curve type that predicts a saturation point (Peat 1970, Paine et al. 2012). Regardless of the specific mechanisms driving increases or decreases in K_h_ over time, applying the asymptotic exponential to predict time-stable K_h_ broadly encompasses all phenomena that may alter K_h_. Our adjustment for K_h_ and K_max_ resulted in VCs showing lower resistance to embolism (Fig. 6), with average *P*_50_ values increasing by 0.33 MPa. Therefore, we suggest to apply this approach to enhance the reliability of VCs, as this makes VCs more suitable for comparison.

*Acer pseudoplatanus*, *Betula pendula*, and *Carpinus betulus* showed in Experiment 1 similar K_h_ trends to those reported for *Corylus avellana*, *Fagus sylvatica*, and *Prunus avium* presented by Silva et al. (2023). However, as expected, the extent of VC shifts (and increases in *P*_50_) varied among the species studied in Experiment 2. While *C. betulus* exhibited the most temporally stable VCs (with *P*_50_ increasing by an average of 4.2%), *F. sylvatica* presented the most unstable VCs over time (with *P*_50_ increasing by an average of 11.3%). Such interspecific variation is a result of species-specific K_h_ dynamics. *C. betulus* demonstrated the most time-stable K_h_ values in Experiment 1, with K_h_ changing by 4.5%. In contrast, *F. sylvatica*, as reported by Silva et al. (2023), exhibited a much more pronounced K_h_ decrease of - 42.3%. The time dependent stability of K_h_ values may be linked to anatomical parameters. *C. betulus*, for instance, has a total pit membrane surface area (*A*_p_) that is at least twice as large as some of the other species studied (see Table S1 and Guan et al. 2022). Anatomical characteristics of the vessels, including vessel diameter, length, and intervessel connectivity, have been associated with the speed of embolism propagation among different vessel-bearing plant species, and consequently linked to the ability of a species to resist loss of conductivity by embolism (Avila et al. 2022b, Pritzkow et al. 2022, Johnson et al. 2023; Lens et al. 2023). However, further investigation involving additional species is necessary to validate this relationship between time-stability of K_h_, anatomical characteristics, and embolism resistance, which lies beyond the scope of this study. It is also unclear to what extent the spin-time artefact may affect flow-centrifuge VCs for gymnosperms.

*P*_50_ values are generally considered as reference point for interspecific comparison of xylem vulnerability to embolism, especially when there are pronounced differences in embolism resistance (Sperry et al. 1988b, Sparks and Black 1999, Domec and Gartner 2001, Sergent et al. 2020). However, when fairly similar or minor differences in embolism resistance occur between samples at the intraspecific or intra-tree level, we argue that including temporal dynamics may improve the accuracy of *P*_50_ estimations, because some species may show a significant increase in *P*_50_ using time-based VCs, while others may show a time-stable *P*_50_. The same suggestion could be applied to *P*_88_ and *P*_12_. Absolute and relative values of *P*_88_ exhibited a similar behaviour to *P*_50_, which is logical because the process influencing *P*_50_ and *P*_88_ is likely similar (i.e., embolism spreading). *P*_12_, however, showed no consistent pattern over time among the samples and species studied, which can be attributed to the varying behaviour of K_h_ values within this Ψ range (between -0.8 and -0.4 MPa).

The most pronounced shifts in VCs occurred after short cumulative spinning times because the most pronounced changes in K_h_ measurements occur during the initial minutes of spinning (Fig. 7 and S4). Long cumulative spinning times tended to improve the sigmoidal shape of VCs, with stable plateaus of high PLC for most of the samples (Fig. 6 and 7). The additional time that is required to generate VCs with a flow-centrifuge following a time-dependent approach, which could be around 30 minutes to ca. 1 hour, appears to be reasonable. Without considerably increasing the overall duration of VC experiments, we suggest acquiring multiple K_h_ measurements within a time frame of 15 minutes at each Ψ step under well-controlled, constant temperature, and then calculating time-stable K_h_ values for VC constructions. This approach enhances the VC accuracy, ensuring K_h_ values are temporally stable, and thus more reliable for comparison between samples.

Although flow-centrifuge represent a highly useful tool for the generation of VCs, the method has not been fully utilized yet in a manner that comprehensively addresses all the physical processes that are involved in embolism spreading (Cochard et al. 2005, Torres-Ruiz et al. 2012, Silva et al. 2023). The data presented show that embolism formation and propagation are not only driven by pressure differences as traditionally assumed (Choat et al. 2010, Torres-Ruiz et al. 2015, Guan et al. 2021, Avila et al. 2022a, Silva et al. 2023), but at least to some extent also affected by a time component. We speculate that gas diffusion kinetics are likely associated with this time-dependent role of embolism propagation, with axial diffusion being considerably faster than radial gas diffusion (Yang et al. 2022, Pereira et al. 2023). Time may play a minor role in other methods that measure embolism resistance, such as other hydraulic methods, as well as the pneumatic, acoustic, and optical approaches. Generally, the non-centrifuge methods rely on bench dehydration of samples, which requires considerable time, and could therefore be less subject to any time related artefact. While this topic requires further research, the approach presented here could also be tested to address time-stability of various parameters in alternative methods.

## Conclusion

Our data confirmed the hypothesis that VCs shift towards higher Ψ values when time-stability of K_h_ is considered in flow-centrifuge measurements. Our analysis indicated the presence of at least two phenomena that may affect K_h_ over time, with one increasing and another decreasing K_h_, and a competitive dynamic between both processes. Although the exact mechanism behind these phenomena remains incompletely understood, its practical implications are relevant: the omission of spin time in flow-centrifuge VCs can introduce an overestimation of embolism resistance, which is on average 8.5% lower in time-stable VCs than VCs that do not account for time-stability of K_h_. The approach presented here may improve the robustness of VC analyses when comparing species, populations, or plants grown under different conditions. Instead of very long spin times to construct VCs, we propose integrating spin time into VCs by acquiring multiple K_h_ measurements at each centrifuge speed for 15 minutes. Moreover, an exponential asymptotic model is recommended to compute time-stable K_h_ values for VCs.

## Data availability statement

The data supporting the findings of this study are available within the paper and supplementary files.

## Acknowledgements

The authors acknowledge Andrea Huppenberger and Clara García Sanchez for technical support. Valuable discussions and insights were provided by Bogusz Bujnowski and Lucian Kaack.

## Author contributions

LMS, LP, MTM and SJ developed the hypotheses and planned the experiments, which were conducted by LMS and JP. LMS analyzed the data. All authors contributed to the interpretation of data and manuscript writing, with substantial inputs from LMS, MTM, LP, and SJ.

## Abbreviations

ΔK_h_: relative change in hydraulic conductivity
*A*_p_: total pit membrane surface area
K_0_: hydraulic conductivity measurements taken immediately after reaching the target Ψ
K_asym_: estimated time- stable K_h_ values derived from the asymptotic exponential model
K_h_: hydraulic conductivity
K_max_: maximum hydraulic conductivity
*L*_V_: average vessel lengths
*P*_12_: *P*_50_ and *P*_88_ water potential values corresponding to 12%, 50%, 88% loss of hydraulic conductivity, respectively
*P*_50___initial_: P_50_ when the cumulative spin time is zero
*P*_50_rel_: the relative value of *P*_50_
PLC: percentage loss of conductivity
RMSE: root mean squared error
VC: vulnerability curve
Ψ: minimum xylem water potential in the middle of a centrifuge sample.

## Supporting Information

**Table S1** Characteristics of the xylem vessels of the three species studied.

**Table S2** Rotational speed (RPM) and water potential (Ψ) applied in experiment 2.

**Fig. S1** Illustration of an asymptotic exponential model curve depicting changes in hydraulic conduc- tivity (K_h_) over time (t).

**Fig. S2** Measured vs. predicted hydraulic conductivity (K_h_) values over time for *A. pseudoplatanus*, *B. pendula*, and *C. betulus* at four water potentials (Ψ).

**Fig. S3** Standardised residual analysis of the asymptotic exponential model from Fig. S2 plot.

**Fig. S4** Vulnerability curves depict the percentage loss of xylem conductivity (PLC) as a function of wa-

ter potential (Ψ) for six angiosperm species.

**Fig. S5** Absolute water potential values corresponding to 88% and 12% loss in conductivity (*P*_88_ and *P*_12_) in (a) and (b), respectively, as a function of cumulative spin time.

**Fig. S6** Relative water potential values corresponding to 88% and 12% loss in conductivity (*P*_88_ and *P*_12_) in (a) and (b), respectively, as a function of cumulative spin time.

**Fig. S7** Slope of the vulnerability curve at water potential values corresponding to 50% loss in conduc- tivity (*S*_50_) as a function of cumulative spin time.

**Fig. S8** Measured (dots) vs. predicted (full red line) hydraulic conductivity (K_h_) values over time for four samples of *Acer pseudoplatanus*, five water potentials (Ψ), and temperatures of 22°C.

**Fig. S9** Measured (dots) vs. predicted (full red line) hydraulic conductivity (K_h_) values over time for four samples of *Betula pendula*, five water potentials (Ψ), and temperatures of 22°C.

**Fig. S10** Measured (dots) vs. predicted (full red line) hydraulic conductivity (K_h_) values over time for four samples of *Carpinus betulus*, seven water potentials (Ψ), and temperatures of 22°C.

**Fig. S11** Measured (dots) vs. predicted (full red line) hydraulic conductivity (K_h_) values over time for four samples of *Corylus avellana*, five water potentials (Ψ), and temperatures of 22°C.

**Fig. S12** Measured (dots) vs. predicted (full red line) hydraulic conductivity (K_h_) values over time for four samples of *Fagus sylvatica*, seven water potentials (Ψ), and temperatures of 22°C.

**Fig. S13** Measured (dots) vs. predicted (full red line) hydraulic conductivity (K_h_) values over time for four samples of *Prunus avium*, five water potentials (Ψ), and temperatures of 22°C.

**Methods S1:** The application of an asymptotic exponential model for estimating time-stable hydraulic conductivity.

## References

1. Alder NN, Pockman WT, Sperry JS, Nuismer S (1997) Use of centrifugal force in the study of xylem cavitation. J Exp Bot 48:665–674.

2. Anderegg WRL, Berry JA, Smith DD, Sperry JS, Anderegg LDL, Field CB (2012) The roles of hydraulic and carbon stress in a widespread climate-induced forest die-off. Proc Natl Acad Sci 109:233–237.

3. Anderegg WRL, Klein T, Bartlett M, Sack L, Pellegrini AFA, Choat B, Jansen S (2016) Meta-analysis reveals that hydraulic traits explain cross-species patterns of drought-induced tree mortality across the globe. Proc Natl Acad Sci 113:5024–5029.

4. Avila RT, Guan X, Kane CN, Cardoso AA, Batz TA, DaMatta FM, Jansen S, McAdam SAM (2022a) Xylem embolism spread is largely prevented by interconduit pit membranes until the majority of conduits are gas-filled. Plant Cell Environ 45:1204–1215.

5. Avila RT, Kane CN, Batz TA, Trabi C, Damatta FM, Jansen S, McAdam SAM (2022b) The relative area of vessels in xylem correlates with stem embolism resistance within and between genera. Tree Physiol 43:75–87.

6. De Baerdemaeker NJF, Arachchige KNR, Zinkernagel J, Van Den Bulcke J, Van Acker J, Schenk HJ, Steppe K, Tognetti R (2019) The stability enigma of hydraulic vulnerability curves: Addressing the link between hydraulic conductivity and drought-induced embolism. Tree Physiol 39:1646–1664.

7. Brodribb TJ, Bowman DJMS, Nichols S, Delzon S, Burlett R (2010) Xylem function and growth rate interact to determine recovery rates after exposure to extreme water deficit. New Phytol 188:533–542.

8. Brodribb T, Brodersen CR, Carriqui M, Tonet V, Rodriguez Dominguez C, McAdam S (2021) Linking xylem network failure with leaf tissue death. New Phytol 232:68–79.

9. Burlett R, Torres-Ruiz JM, Tyree M, Delzon S, Cochard H (2022) Establishing vulnerability curves with the flow-through centrifuge method (cavitron) Protocol. :1–17. https://prometheusprotocols.net/function/water-relations/xylem-vulnerability-%20curves/establishing-vulnerability-curves-with-the-flow-through-centrifuge-method-cavitron/

10. Cai J, Tyree MT (2010) The impact of vessel size on vulnerability curves: Data and models for within- species variability in saplings of aspen, Populus tremuloides Michx. Plant Cell Environ 33:1059– 1069.

11. Canny MJ, Sparks JP, Huang CX, Roderick ML (2007) Air embolisms exsolving in the transpiration water - The effect of constrictions in the xylem pipes. Funct Plant Biol 34:95–111.

12. Choat B, Drayton WM, Brodersen C, Matthews MA, Shackel KA, Wada HIR, McElrone AJ (2010) Measurement of vulnerability to water stress-induced cavitation in grapevine: A comparison of four techniques applied to a long-vesseled species. Plant Cell Environ 33:1502–1512.

13. Choat B, Jansen S, Brodribb TJ, Cochard H, Delzon S, Bhaskar R, Bucci SJ, Feild TS, Gleason SM, Hacke UG, Jacobsen AL, Lens F, Maherali H, Martínez-Vilalta J, Mayr S, Mencuccini M, Mitchell PJ, Nardini A, Pittermann J, Pratt RB, Sperry JS, Westoby M, Wright IJ, Zanne AE (2012) Global convergence in the vulnerability of forests to drought. Nature 491:752–755.

14. Cochard H (1992) Vulnerability of several conifers to air embolism. Tree Physiol 11: 73–83.

15. Cochard H (2002) A technique for measuring xylem hydraulic conductance under high negative pressures. Plant Cell Environ 25:815–819. https://onlinelibrary.wiley.com/terms-and-conditions

16. Cochard H, Badel E, Herbette S, Delzon S, Choat B, Jansen S (2013) Methods for measuring plant vulnerability to cavitation: A critical review. J Exp Bot 64:4779–4791.

17. Cochard H, Barigah T, Herbert E, Caupin F (2007) Cavitation in plants at low temperature: Is sap transport limited by the tensile strength of water as expected from Briggs’ Z-tube experiment? New Phytol 173:571–575.

18. Cochard H, Bodet C, Améglio T, Cruiziat P (2000) Cryo-scanning electron microscopy observations of vessel content during transpiration in walnut petioles. Facts or artefacts? Plant Physiol 124:1191– 1202.

19. Cochard H, Damour G, Bodet C, Tharwat I, Poirier M, Améglio T (2005) Evaluation of a new centrifuge technique for rapid generation of xylem vulnerability curves. Physiol Plant 124:410–418.

20. Domec JC, Gartner BL (2001) Cavitation and water storage capacity in bole xylem segments of mature and young Douglas-fir trees. Trees 15:204–214.

21. Duursma RA, Choat B (2016) fitplc-an R package to fit hydraulic vulnerability curves. J Plant Hydraul 4: e002

22. Espino S, Schenk HJ (2011) Mind the bubbles: Achieving stable measurements of maximum hydraulic conductivity through woody plant samples. J Exp Bot 62:1119–1132.

23. Goudriaan J (1979) A family of saturation type curves, especially in relation to photosynthesis. Ann Bot 43:783–785.

24. Guan X, Pereira L, McAdam SAM, Cao KF, Jansen S (2021) No gas source, no problem: Proximity to pre- existing embolism and segmentation affect embolism spreading in angiosperm xylem by gas diffusion. Plant Cell Environ 44:1329–1345.

25. Guan X, Werner J, Cao KF, Pereira L, Kaack L, McAdam SAM, Jansen S (2022) Stem and leaf xylem of angiosperm trees experiences minimal embolism in temperate forests during two consecutive summers with moderate drought. Plant Biol 24:1208–1223.

26. Hacke UG, Sperry JS (2001) Functional and ecological xylem anatomy. Perspect Plant Ecol Evol Syst 4:97–115.

27. Holbrook NM, Burns MJ, Field CB (1995) Negative Xylem Pressures in Plants: A Test of the Balancing Pressure Technique. Sci 270:1193–1194.

28. Jansen S, Gortan E, Lens F, Lo Gullo MA, Salleo S, Scholz A, Stein A, Trifilò P, Nardini A (2011) Do quantitative vessel and pit characters account for ion-mediated changes in the hydraulic conductance of angiosperm xylem? New Phytol 189:218–228.

29. Jansen S, Schenk HJ (2015) On the ascent of sap in the presence of bubbles. Am J Bot 102:1561–1563.

30. Johnson KM, Everbach SR, Holbrook NM, Olson ME (2023) Evaluating Carlquist’s Law from a physiological perspective. IAWA J 45:1–10.

31. Krieger L, Schymanski SJ (2023) A new experimental setup to measure hydraulic conductivity of plant segments. AoB Plants 15:1–13.

32. Lens F, Gleason SM, Bortolami G, Brodersen C, Delzon S, Jansen S (2022) Functional xylem characteristics associated with drought-induced embolism in angiosperms. New Phytol 236:2019–2036.

33. Lintunen A, Salmon Y, Hölttä T, Suhonen H (2022) Inspection of gas bubbles in frozen Betula pendula xylem with micro-CT: Conduit size, water status and bark permeability affect bubble characteristics. Physiol Plant 174:1–11.

34. López R, Nolf M, Duursma RA, Badel E, Flavel RJ, Cochard H, Choat B (2018) Mitigating the open vessel artefact in centrifuge-based measurement of embolism resistance. Tree Physiol 39:143–155.

35. Martin-StPaul NK, Longepierre D, Huc R, Delzon S, Burlett R, Joffre R, Rambal S, Cochard H (2014) How reliable are methods to assess xylem vulnerability to cavitation? The issue of ‘open vessel’ artifact in oaks. Tree Physiol 34:894–905.

36. Miguez F, Archontoulis S, Dokoohaki H (2018) Chapter 15: Nonlinear Regression Models and Applications. In: Barry G, Kathleen MY, editors. Applied Statistics in Agricultural, Biological, and Environmental Sciences, Madison, WI.

37. Neufeld HS, Grantz DA, Meinzer FC, Goldstein G, Crisosto GM, Crisosto C (1992) Genotypic variability in vulnerability of leaf xylem to cavitation in water-stressed and well-irrigated sugarcane’. Plant Physiol 100:1020–1028. https://academic.oup.com/plphys/article/100/2/1020/6085900

38. Paine CET, Marthews TR, Vogt DR, Purves D, Rees M, Hector A, Turnbull LA (2012) How to fit nonlinear plant growth models and calculate growth rates: An update for ecologists. Methods Ecol Evol 3:245–256.

39. Pammenter NW, Van der Willigen C (1998) A mathematical and statistical analysis of the curves illustrating vulnerability of xylem to cavitation. Tree Physiol 18:589–593.

40. Peat WE (1970) Relationships between Photosynthesis and Light Intensity in the Tomato. Ann Bot 34:319–328.

41. Pereira L, Kaack L, Guan X, Silva L de M, Miranda MT, Pires GS, Ribeiro R V., Schenk HJ, Jansen S (2023) Angiosperms follow a convex trade-off to optimize hydraulic safety and efficiency. New Phytol 240:1788–1801.

42. Pockman WT, Sperry JS, O’leary JW (1995) Sustained and significant negative water pressure in xylem. Nature 378:715–716.

43. Pritzkow C, Brown MJM, Carins-Murphy MR, Bourbia I, Mitchell PJ, Brodersen C, Choat B, Brodribb TJ (2022) Conduit position and connectivity affect the likelihood of xylem embolism during natural drought in evergreen woodland species. Ann Bot 130:431–444.

44. Scholz A, Klepsch M, Karimi Z, Jansen S (2013) How to quantify conduits in wood? Front Plant Sci 4:1– 12.

45. Sergent AS, Varela SA, Barigah TS, Badel E, Cochard H, Dalla-Salda G, Delzon S, Fernández ME, Guillemot J, Gyenge J, Lamarque LJ, Martinez-Meier A, Rozenberg P, Torres-Ruiz JM, Martin- StPaul NK (2020) A comparison of five methods to assess embolism resistance in trees. For Ecol Manage 468

46. Silva LM, Pereira L, Kaack L, Guana X, Trabia CL, Jansen S (2023) Gas diffusion kinetics drive embolism spread in angiosperm xylem: evidence from flow- centrifuge experiments and modelling. bioRxiv: doi: 10.1101/2023.04.19.537442.

47. Silva LM, Pereira L, Kaack L, Guana X, Trabia CL, Jansen S (submitted) Gas diffusion kinetics drive embolism spread in angiosperm xylem: evidence from flow- centrifuge experiments and modelling. New Phytol

48. Sorz J, Hietz P (2006) Gas diffusion through wood: Implications for oxygen supply. Trees 20:34–41.

49. Sparks JP, Black RA (1999) Regulation of water loss in populations of Populus trichocarpa: the role of stomatal control in preventing xylem cavitation. Tree Physiol 19:453–459.

50. Sperry JS (1985) Xylem embolism in the Palm Rhapis excelsa. IAWA J 6:283–292.

51. Sperry JS, Donnelly JR, Tyree MT (1988a) A method for measuring hydraulic conductivity and embolism in xylem. Plant Cell Environ 11:35–40.

52. Sperry JS, Tyree MT, Donnelly JR (1988b) Vulnerability of xylem to embolism in a mangrove vs an inland species of Rhizophoraceae. Physiol Plant 74:276–283.

53. Taneda H, Sperry JS (2008) A case-study of water transport in co-occurring ring- versus diffuse-porous trees: contrasts in water-status, conducting capacity, cavitation and vessel refilling. Tree Physiol 28:1641–1651. http://treephys.oxfordjournals.org/

54. Torres-Ruiz JM, Jansen S, Choat B, McElrone AJ, Cochard H, Brodribb TJ, Badel E, Burlett R, Bouche PS, Brodersen CR, Li S, Morris H, Delzon S (2015) Direct X-ray microtomography observation confirms the induction of embolism upon xylem cutting under tension. Plant Physiol 167:40–43.

55. Torres-Ruiz JM, Sperry JS, Fernández JE (2012) Improving xylem hydraulic conductivity measurements by correcting the error caused by passive water uptake. Physiol Plant 146:129–135.

56. Tyree MT, Fiscus EL, Wullschleger SD, Dixon MA (1986) Detection of Xylem Cavitation in Corn under Field Conditions. Plant Physiol. 82: 597–599

57. Tyree MT, Patiño S, Becker P (1998) Vulnerability to drought-induced embolism of Bornean heath and dipterocarp forest trees. Tree Physiol 18:583–588.

58. Tyree MT, Sperry JS (1988) Do Woody Plants Operate Near the Point of Catastrophic Xylem Dysfunction Caused by Dynamic Water Stress? Plant Physiol 88:574–580.

59. Tyree MT, Sperry JS (1989) Vulnerability of Xylem to Cavitation and Embolism. Annu Rev Plant Physiol Plant Mol Biol 40:19–36.

60. Tyree MT, Zimmermann MH (2002) Xylem Structure and the Ascent of Sap. In: Timell TE, editor. 2nd ed. Springer-Verlag Berlin Heidelberg, Syracuse.

61. Wang Y, Burlett R, Feng F, Tyree MT (2014a) Improved precision of hydraulic conductance measurements using a Cochard rotor in two different centrifuges. J Plant Hydraul 1:e007.

62. Wang Y, Pan R, Tyree MT (2015) Studies on the tempo of bubble formation in recently cavitated vessels: A model to predict the pressure of air bubbles. Plant Physiol 168:521–531.

63. Wang R, Zhang L, Zhang S, Cai J, Tyree MT (2014b) Water relations of Robinia pseudoacacia L.: Do vessels cavitate and refill diurnally or are R-shaped curves invalid in Robinia? Plant Cell Environ 37:2667–2678.

64. Wheeler JK, Huggett BA, Tofte AN, Rockwell FE, Holbrook NM (2013) Cutting xylem under tension or supersaturated with gas can generate PLC and the appearance of rapid recovery from embolism. Plant Cell Environ 36:1938–1949.

65. Yang D, Pereira L, Peng G, Ribeiro R V, Kaack L, Tyree MT, Jansen S, Tyree MT (2022) A Unit Pipe Pneumatic model to simulate gas kinetics during measurements of embolism in 2 excised angiosperm xylem Corresponding Authors. Tree Physiol 43:88–101.

66. Yang S, Tyree MT (1992) A theoretical model of hydraulic conductivity recovery from embolism with comparison to experimental data on Acer saccharum. Plant Cell Environ 15:633–643.

67. Yin P, Meng F, Liu Q, An R, Cai J, Du G (2019) A comparison of two centrifuge techniques for constructing vulnerability curves: insight into the ‘open-vessel’ artifact. Physiol Plant 165:701– 710.

68. Zhang Y, Carmesin C, Kaack L, Klepsch MM, Kotowska M, Matei T, Schenk HJ, Weber M, Walther P, Schmidt V, Jansen S (2020) High porosity with tiny pore constrictions and unbending pathways characterize the 3D structure of intervessel pit membranes in angiosperm xylem. Plant Cell Environ 43:116–130.

69. Zhang Y, Pereira L, Kaack L, Liu J, Jansen S (2024) Gold perfusion experiments support the multi-layered, mesoporous nature of intervessel pit membranes in angiosperm xylem. New Phytol 242:493– 506.

