## Supplementary material for "Time-based shifts in xylem vulnerability curves of angiosperms based on the flow-centrifuge method": Fig. S

The following Supporting Information is available for this article:

**Table S1** Characteristics of the xylem vessels of the three species studied.

**Table S2** Rotational speed (RPM) and water potential (Ψ) applied in experiment 2.

**Table S1** Characteristics of the xylem vessels of the six species studied. Values indicate the mean and standard deviation of the equivalent circle diameter of vessels (*D*), mean vessel length (*L_v_*), maximum vessel length (*L_v_max_*), thickness of an intervessel pit membrane at the centre (*T*_PM_), total pit membrane surface area (*A_p_*), and vessel volume (*V_v_*). Values of *D*, *L_v_*, *L_v_max_*, and *T*_PM_ were obtained from Guan et al. (2022). The estimation of *A_p_* followed the model of Wheeler et al. (2005). *V*_v_ was calculated by considering a vessel as a cylinder, with *V_v_* = π (*D*/2)^2^ × *L_v_*.

| Species | *D* (µm) | *L*_V_ (cm) | *L*_V_max_ (cm) | *T*_PM_ (nm) | *A_p_* (µm^2^) | | *V_v_* (10^-11^ m^3^) | |
| --- | --- | --- | --- | --- | --- | --- | --- | --- |
| *Acer pseudoplatanus* | 29.56±12.33 | 3.98±1.62 | 11.78±2.65 | 360±574 | | 0.552 | | 2.74×10^-11^ |
| *Betula pendula* | 20.08±7.05 | 5.73±1.35 | 13.64±0.72 | 191±39 | | 0.382 | | 1.81×10^-11^ |
| *Carpinus betulus* | 20.53±10.07 | 5.50±0.70 | 12.50±4.55 | 296±58 | | 1.045 | | 1.82×10^-11^ |
| *Corylus avellana* | 19.78±7.17 | 3.55±0.83 | 11.6±1.36 | 251±35 | | 0.314 | | 1.09×10^-11^ |
| *Fagus sylvatica* | 16.58±4.96 | 5.15±0.82 | 13.58±3.54 | 244±58 | | 0.118 | | 1.11×10^-11^ |
| *Prunus avium* | 16.02±5.08 | 10.6±1.67 | 22.30±1.76 | 385±86 | | 0.470 | | 2.14×10^-11^ |

**Table S2** Rotational speed (RPM) and water potential (Ψ) applied in experiment 2 for constructing vulnerability curves with a flow-centrifuge method for six angiosperm species.

| *Acer pseudoplatanus* | | *Betula pendula* | | *Carpinus betulus* | | *Corylus avellana* | | *Fagus sylvatica* | | *Prunus avium* | |
| --- | --- | --- | --- | --- | --- | --- | --- | --- | --- | --- | --- |
| RPM | Ψ (MPa) | RPM | Ψ (MPa) | RPM | Ψ (MPa) | RPM | Ψ (MPa) | RPM | Ψ (MPa) | RPM | Ψ (MPa) |
| 2000 | -0.35 | 2000 | -0.35 | 2000 | -0.35 | 2000 | -0.35 | 2000 | -0.35 | 2000 | -0.35 |
| 3000 | -0.79 | 3000 | -0.79 | 3000 | -0.79 | 3000 | -0.79 | 3000 | -0.79 | 3000 | -0.79 |
| 5800 | -2.97 | 4000 | -1.41 | 5000 | -2.21 | 4250 | -1.59 | 4875 | -2.10 | 5800 | -2.97 |
| 6400 | -3.62 | 4200 | -1.56 | 5700 | -2.87 | 4875 | -2.10 | 5500 | -2.67 | 6600 | -3.84 |
| 7000 | -4.32 | 4500 | -1.79 | 6000 | -3.18 | 5250 | -2.43 | 6250 | -3.45 | 7100 | -4.45 |
|  | - | - | - | 6500 | -3.73 |  | - |  |  |  | - |

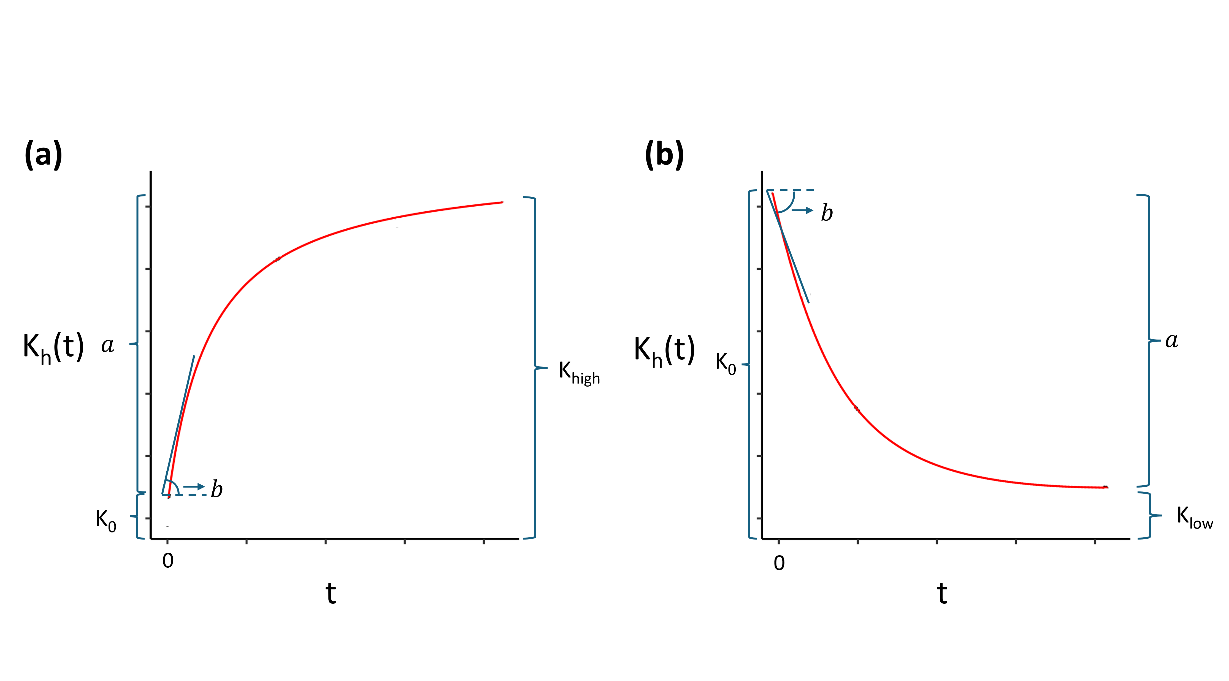

**Fig. S1** Illustration of an asymptotic exponential model curve depicting changes in hydraulic conductivity (K_h_) over time (t). In this model, $a$ denotes the asymptote coefficient, *b* represents the initial slope of the curve at the beginning of the time series, and K_0_ is the intercept denoting the hydraulic conductivity measured at t=0. Panel (a) depicts an instance of K_h_ increasing over time, estimating the highest hydraulic conductivity values (K_high_), while panel (b) illustrates K_h_ decreasing over time, estimating the lowest value of hydraulic conductivity (K_low_).

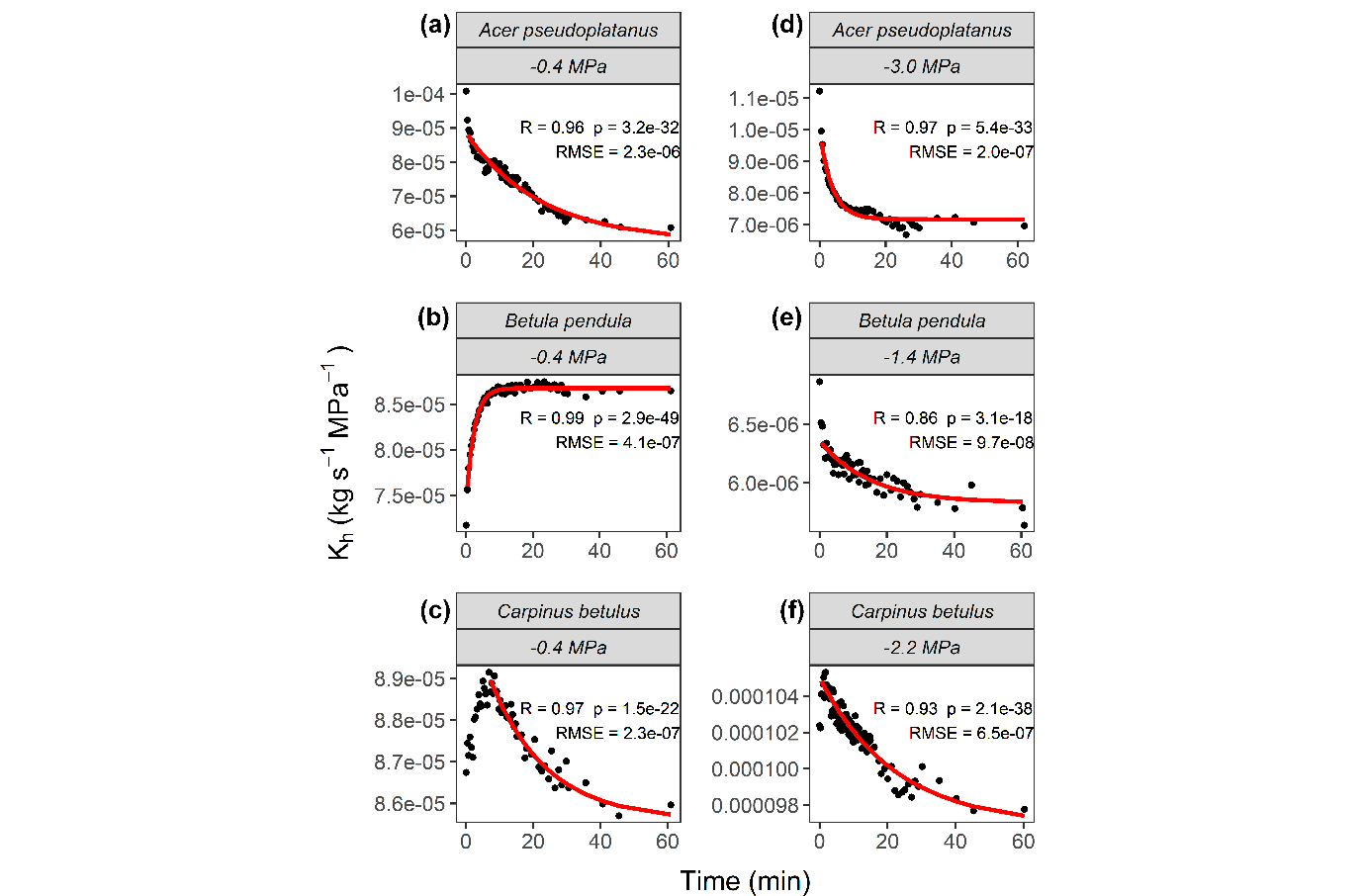

**Fig. S2** Measured (dots) vs. predicted (full red line) hydraulic conductivity (K_h_) values over time for three angiosperm species (*A. pseudoplatanus*, *B. pendula*, and *C. betulus*) at four water potentials (Ψ) and temperatures of 22^o^C. Dots represent stem samples that were spun in a flow-centrifuge for 1 hour at a constant temperature and subjected to a constant Ψ based on a given rotational speed. The curves were predicted values of K_h_, which were estimated by an asymptotic exponential model. The root mean squared error (RMSE), R-squared (R^2^), and p-value (p) were used as a measure of the goodness of fit.

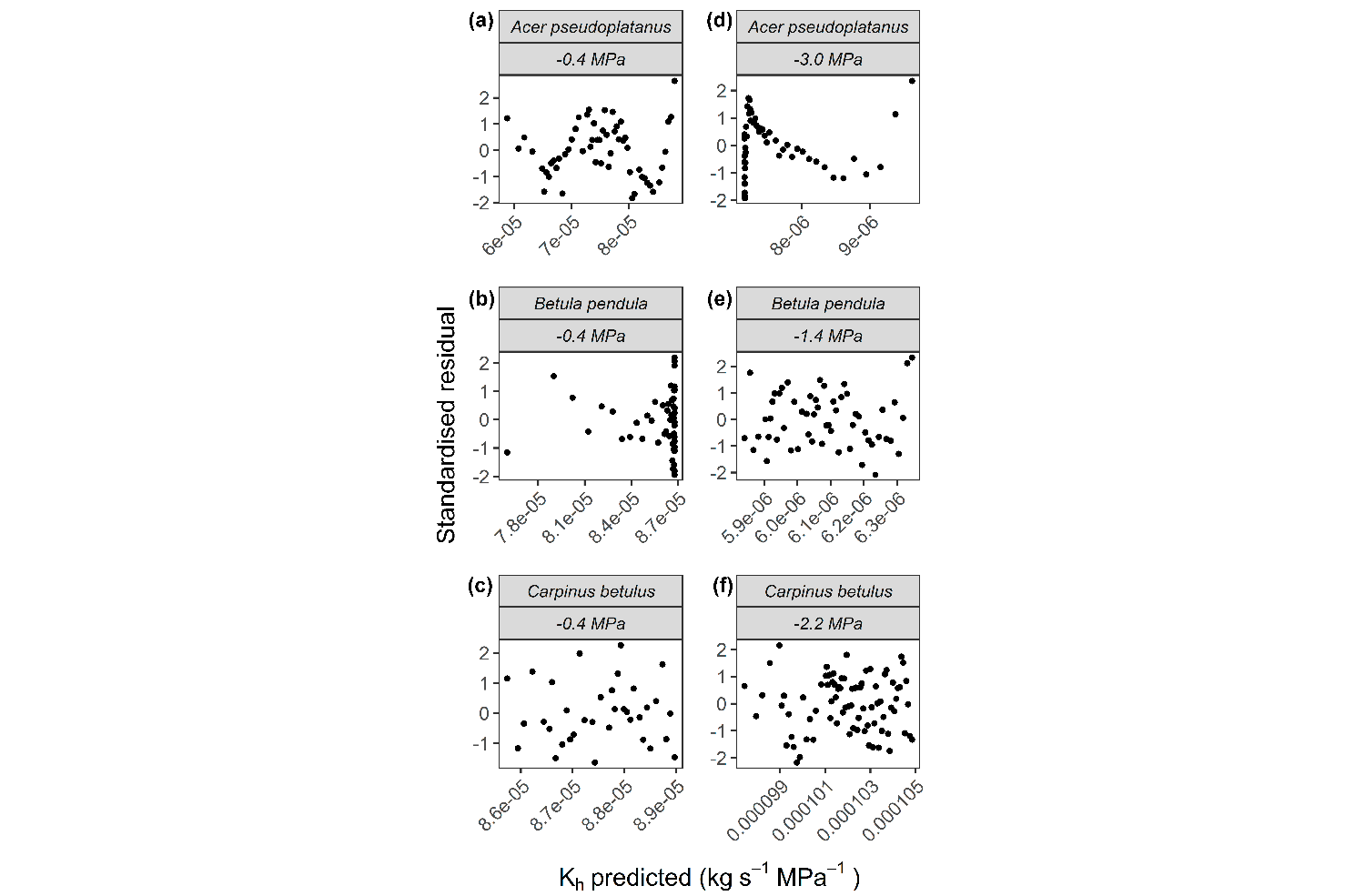

**Fig. S3** Standardised residual analysis of the asymptotic exponential model (Eq. 6 and 8) from Fig. S2, depicting the relationship between predicted values of hydraulic conductivity (K_h_) and residuals for three angiosperm species (*A. pseudoplatanus*, *B. pendula*, and *C. betulus*) at four water potentials (Ψ) and temperatures of 22^o^C.

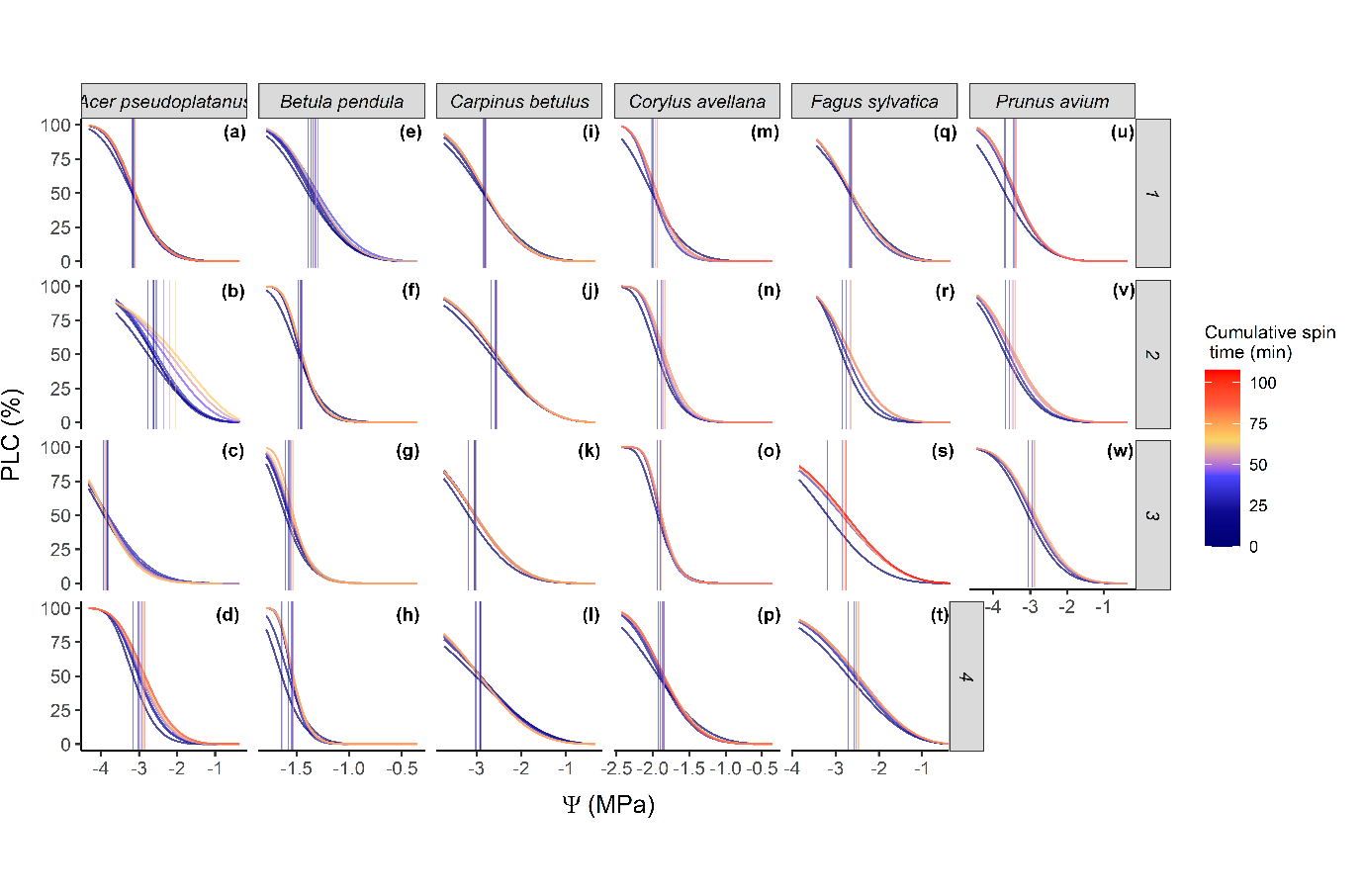
 **Fig. S4** Vulnerability curves depict the percentage loss of xylem hydraulic conductivity (PLC) as a function of water potential (Ψ) for six angiosperm species. These curves were generated from data depicted in Fig 2 by grouping measurements based on the time elapsed since reaching the target Ψ in a flow-centrifuge. The figure presents a grid layout with species (columns) and samples (rows). Each sample of *Acer pseudoplatanus*, *Betula pendula*, and *Carpinus betulus*, includes 7 vulnerability curves, while each sample of *Corylus avellana*, *Fagus sylvatica*, and *Prunus avium* includes 4 vulnerability curves. Each curve is distinguished by a different colour, representing cumulative spin time, which is calculated as the accumulated time spent at each pre-set centrifuge speed. Vertical line represents the Ψ corresponding to 50% loss in conductivity (*P*_50_), which were also coloured according to the cumulative spin time.

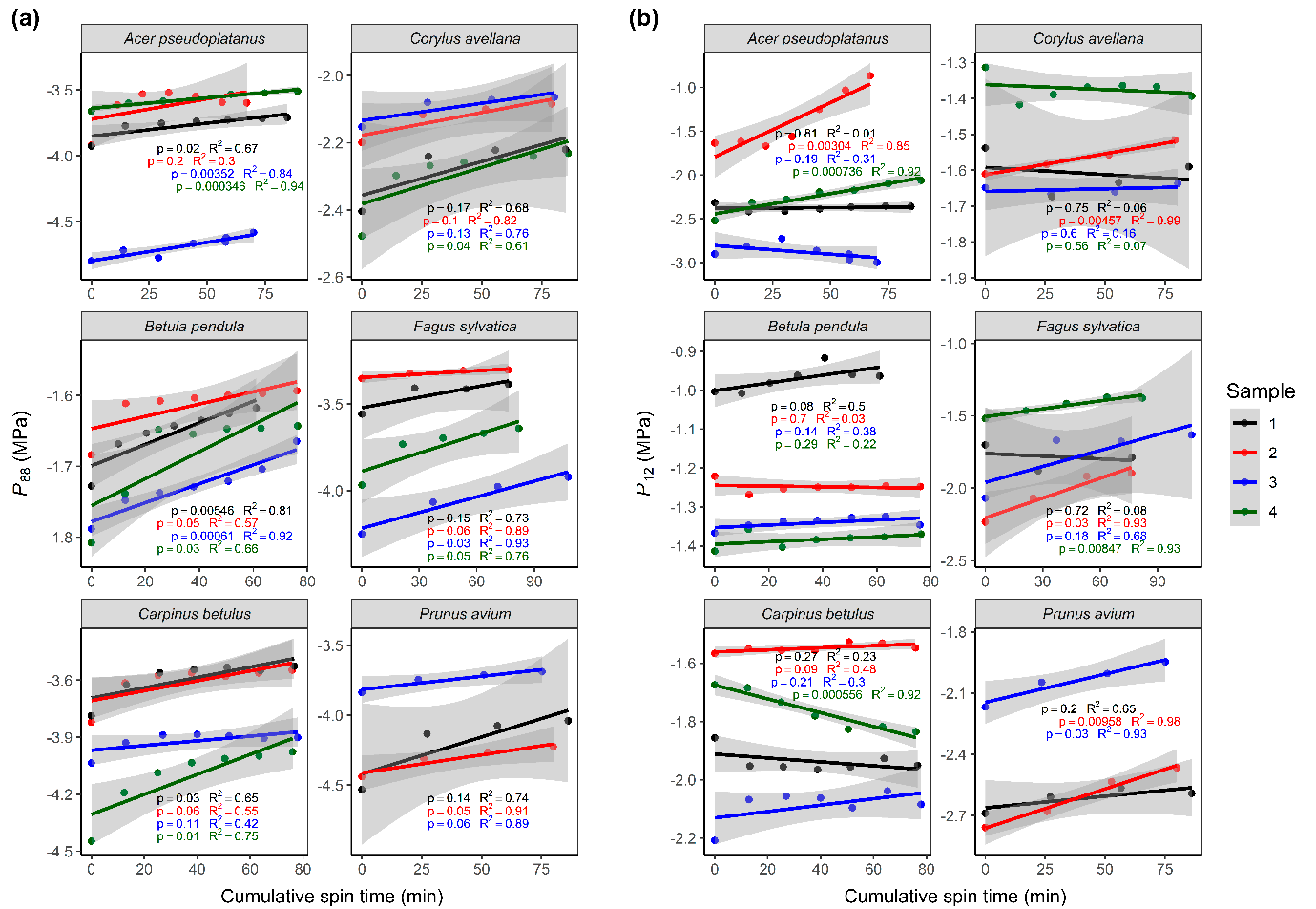

**Fig. S5** Absolute water potential values corresponding to 88% and 12% loss in conductivity (*P*_88_ and *P*_12_) in (a) and (b), respectively, as a function of cumulative spin time for six angiosperm species. Cumulative spin time is calculated as the accumulated time spent at each pre-set centrifuge speed. Each colour represents a different sample, with n = 4 for *Acer pseudoplatanus*, *Betula pendula*, *Carpinus betulus*, *Corylus avellana*, and *Fagus sylvatica*, and n = 3 for *Prunus avium*. Points represent estimated *P*_88_ or *P*_12_ values obtained from vulnerability curves in Fig S4. Coloured lines indicate the correlation between *P*_88_ or *P*_12_ and cumulative spin time, with corresponding confidence intervals in grey, and p-value (p) and R-squared (R^2^) values provided.

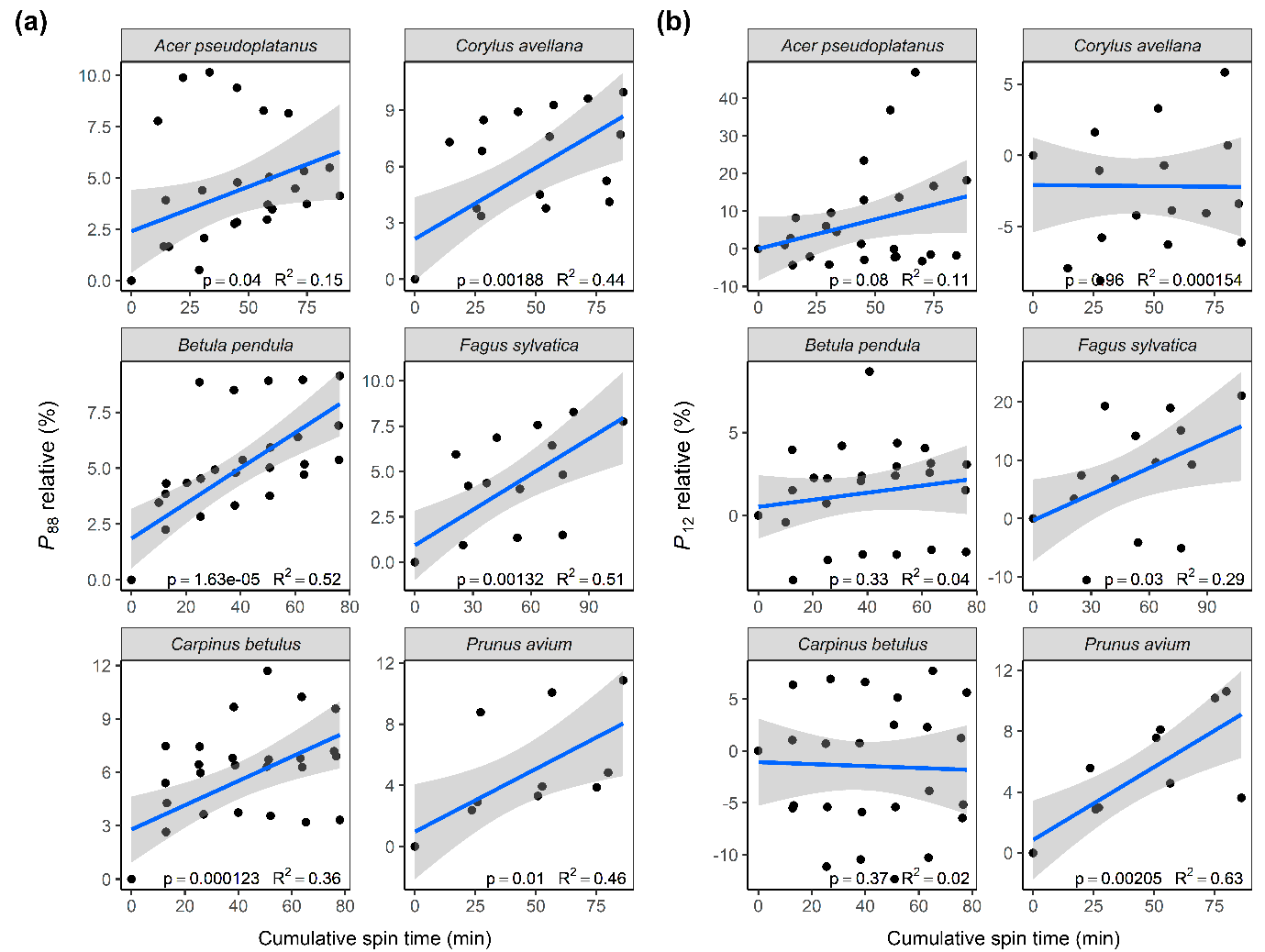

**Fig. S6** Relative water potential values corresponding to 88% and 12% loss in conductivity (*P*_88_ and *P*_12_) in (a) and (b), respectively, as a function of cumulative spin time for six angiosperm species (*Acer pseudoplatanus*, *Betula pendula*, *Carpinus betulus*, *Corylus avellana*, *Fagus sylvatica*, and *Prunus avium*). Cumulative spin time is calculated as the accumulated time spent at each pre-set centrifuge speed. Points represent the relative values of *P*_88_ or *P*_12_ that were calculated through equation 3 and represent the temporal changes relative to their value when cumulative spin time was equal to zero. The blue line indicates the correlation between *P*_88_ or *P*_12_ relative and cumulative spin time, with corresponding confidence intervals in grey, and p-value (p) and R-squared (R^2^) values provided.

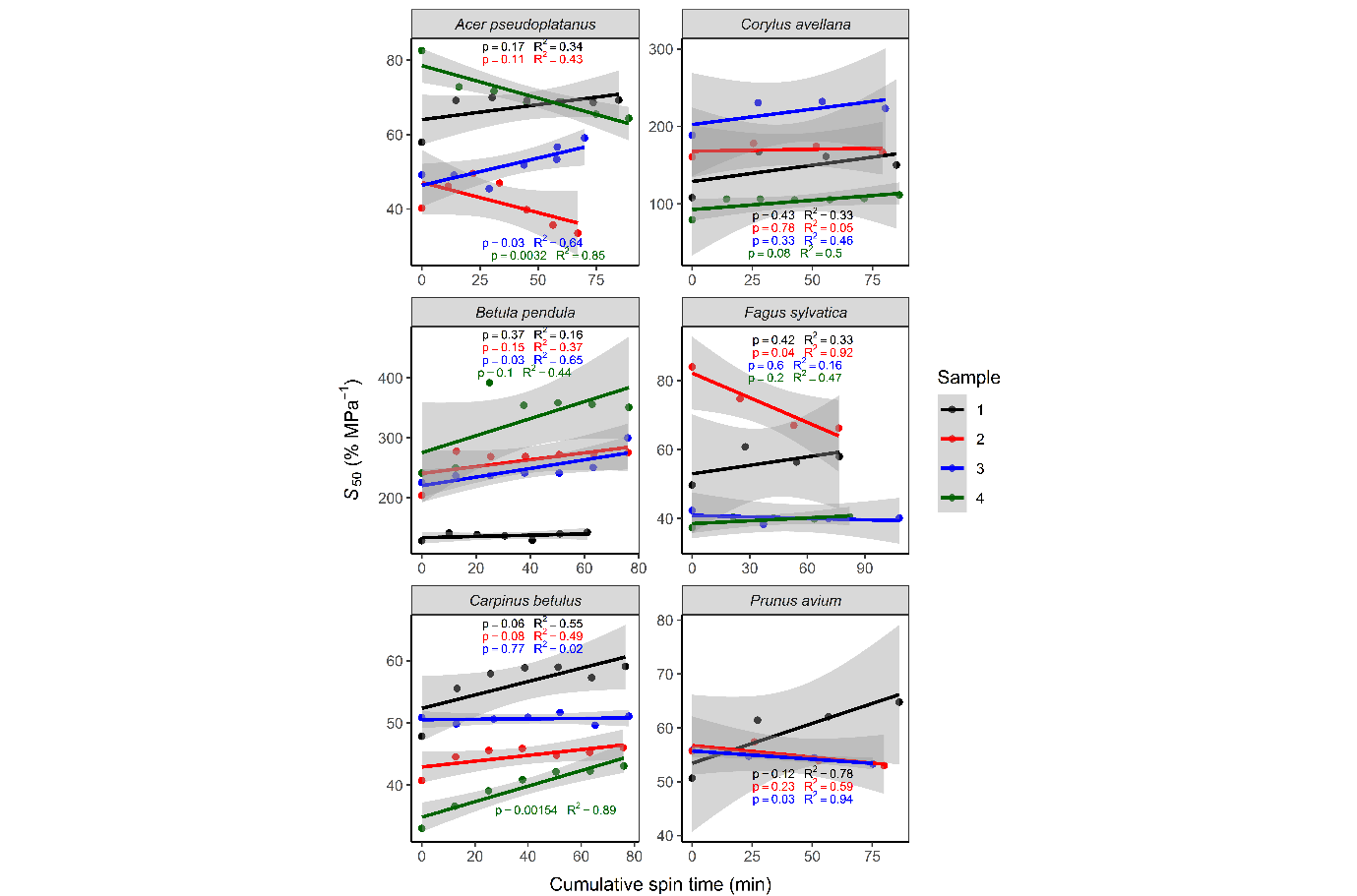

**Fig. S7** Slope of the vulnerability curve at water potential values corresponding to 50% loss in conductivity (*S*_50_) as a function of cumulative spin time for six angiosperm species. Cumulative spin time is calculated as the accumulated time spent at each pre-set centrifuge speed. Each colour represents a different sample, with n = 4 for *Acer pseudoplatanus*, *Betula pendula*, *Carpinus betulus*, *Corylus avellana*, and *Fagus sylvatica*, and n = 3 for *Prunus avium*. Points represent estimated *S*_50_ values obtained from vulnerability curves in Fig S4. Coloured lines indicate the correlation between *S*_50_ and cumulative spin time, with corresponding confidence intervals in grey, and p-value (p) and R-squared (R^2^) values provided.

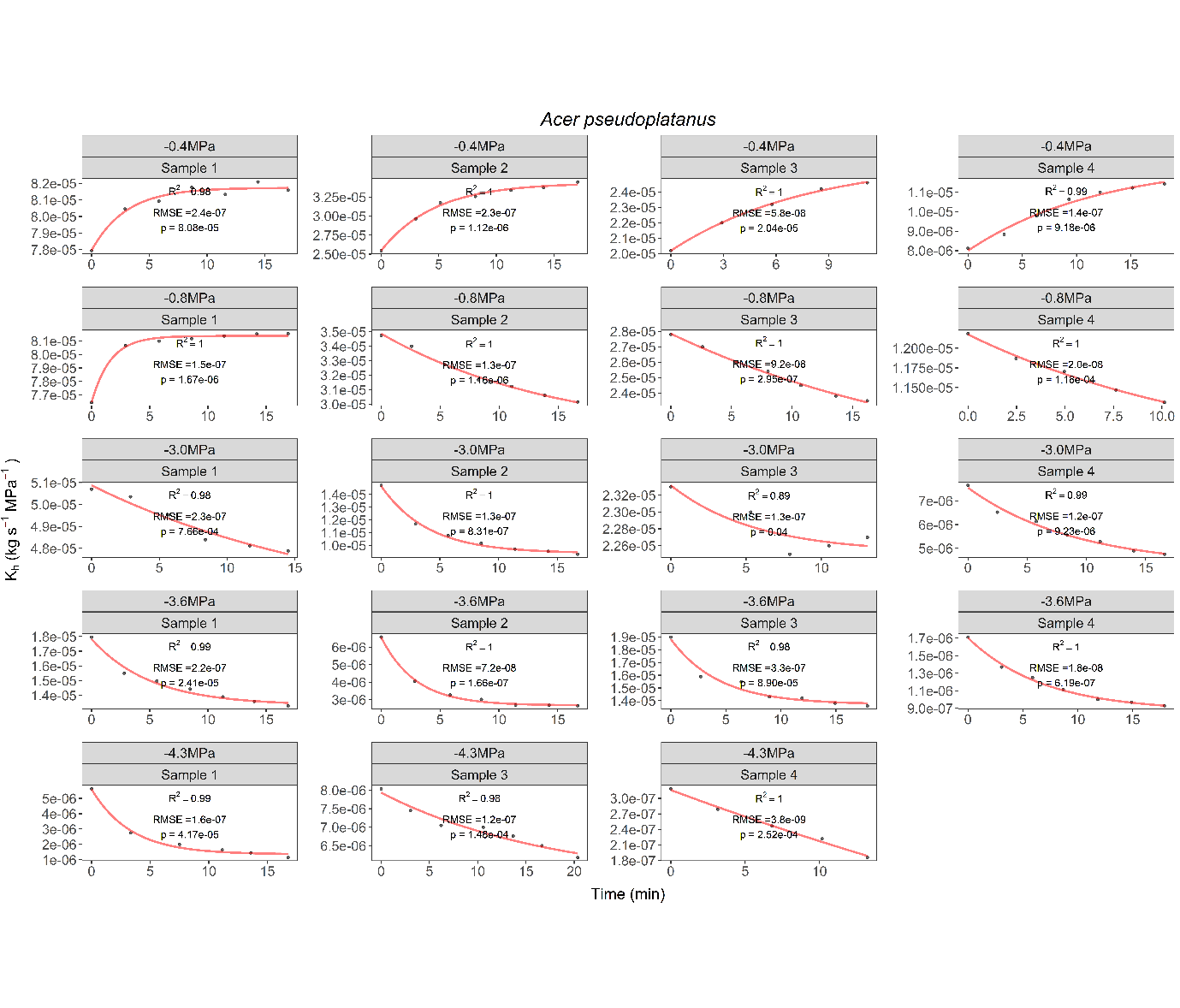

**Fig. S8** Measured (dots) vs. predicted (full red line) hydraulic conductivity (K_h_) values over time for four samples of *Acer pseudoplatanus*, five water potentials (Ψ), and temperatures of 22^o^C. Dots represent stem samples that were spun in a flow-centrifuge for 15 minutes at a constant temperature and subjected to a constant Ψ based on a given rotational speed. The curves were predicted values of K_h_, which were estimated by an asymptotic exponential model. The root mean squared error (RMSE), R-squared (R^2^), and p-values (p) were used as a measure of the goodness of fit.

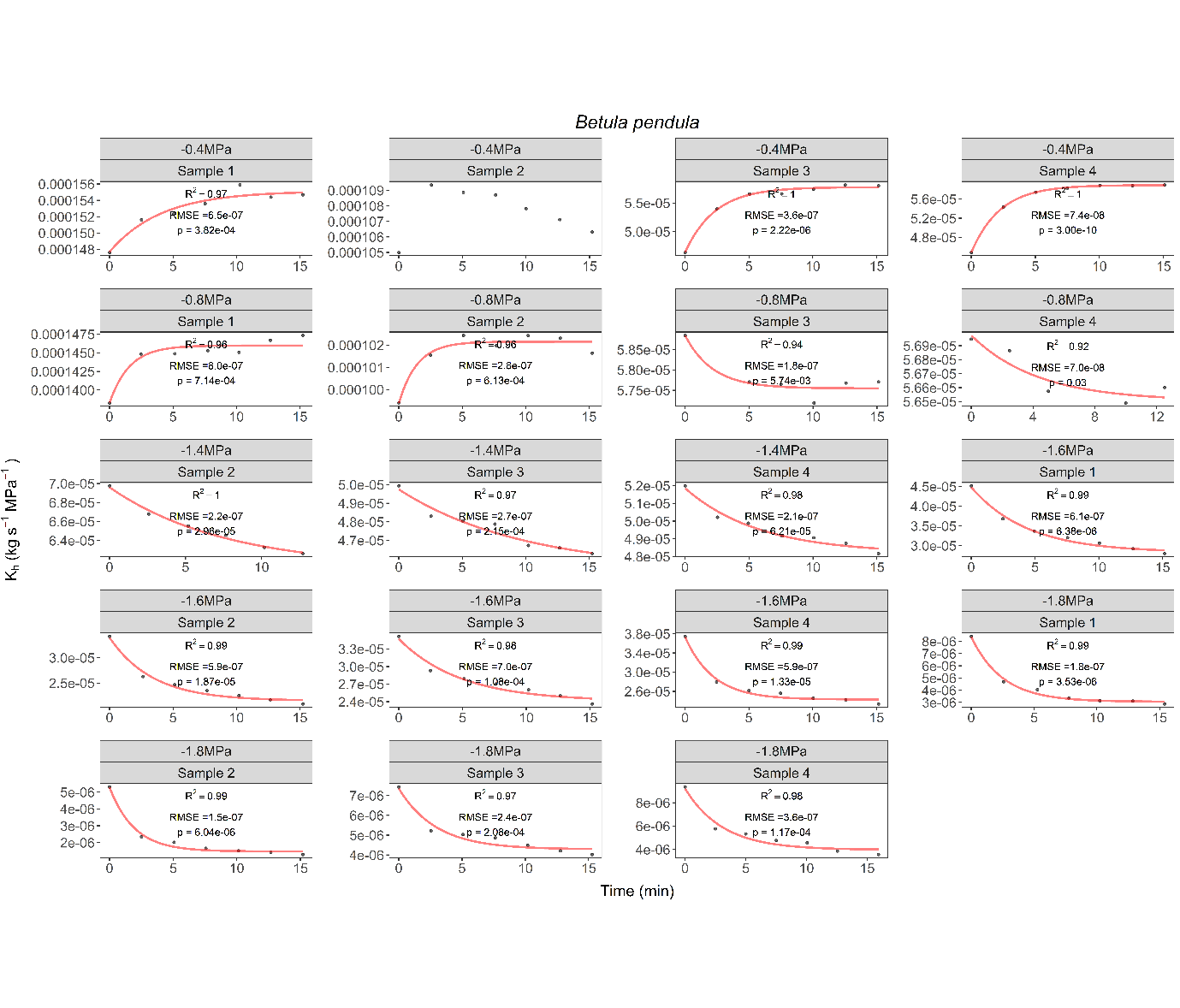

**Fig. S9** Measured (dots) vs. predicted (full red line) hydraulic conductivity (K_h_) values over time for four samples of *Betula pendula*, five water potentials (Ψ), and temperatures of 22^o^C. Dots represent stem samples that were spun in a flow-centrifuge for 15 minutes at a constant temperature and subjected to a constant Ψ based on a given rotational speed. The curves were predicted values of K_h_, which were estimated by an asymptotic exponential model. The root mean squared error (RMSE), R-squared (R^2^), and p-values (p) were used as a measure of the goodness of fit.

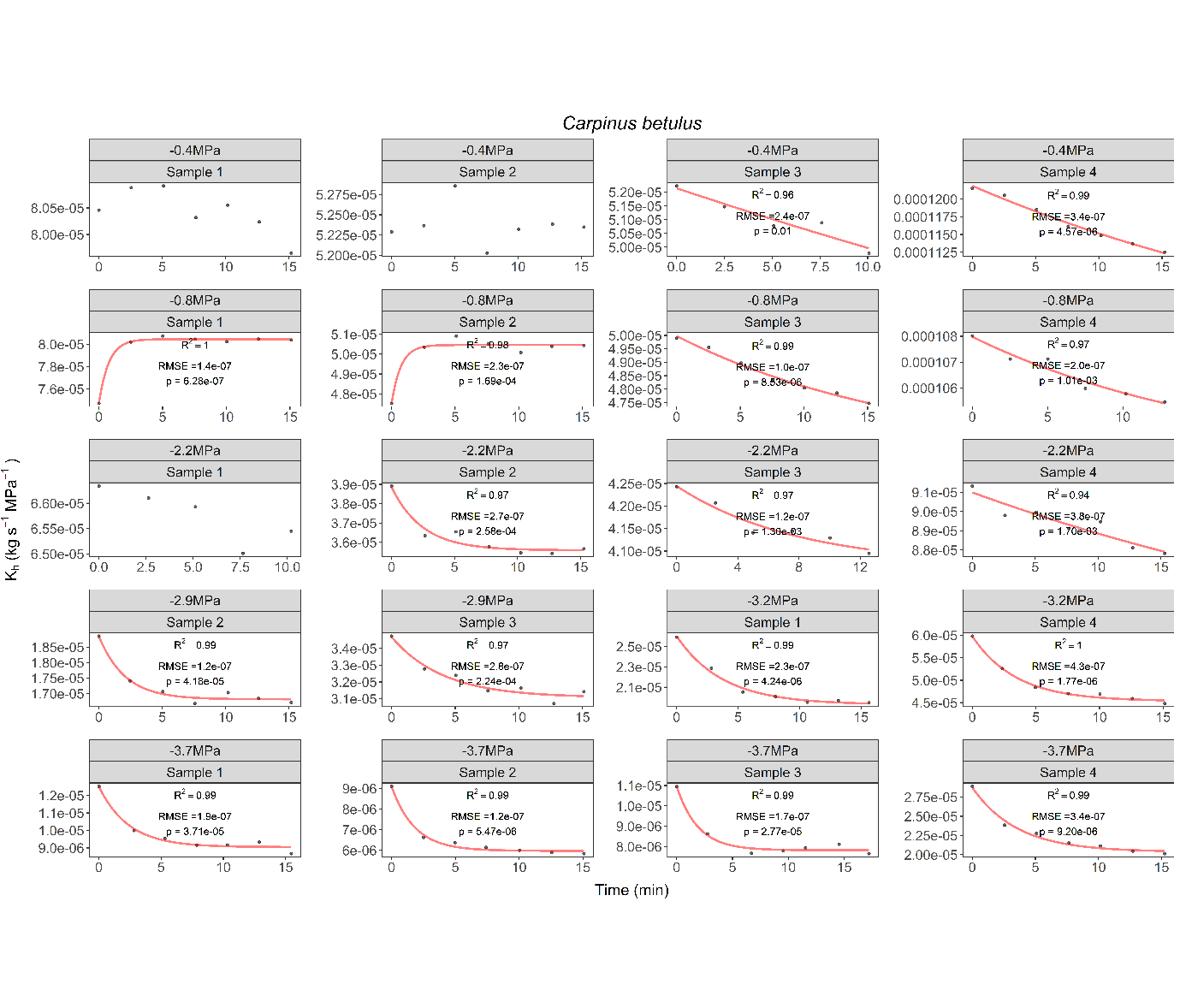

**Fig. S10** Measured (dots) vs. predicted (full red line) hydraulic conductivity (K_h_) values over time for four samples of *Carpinus betulus*, seven water potentials (Ψ), and temperatures of 22^o^C. Dots represent stem samples that were spun in a flow-centrifuge for 15 minutes at a constant temperature and subjected to a constant Ψ based on a given rotational speed. The curves were predicted values of K_h_, which were estimated by an asymptotic exponential model. The root mean squared error (RMSE), R-squared (R^2^), and p-values (p) were used as a measure of the goodness of fit.

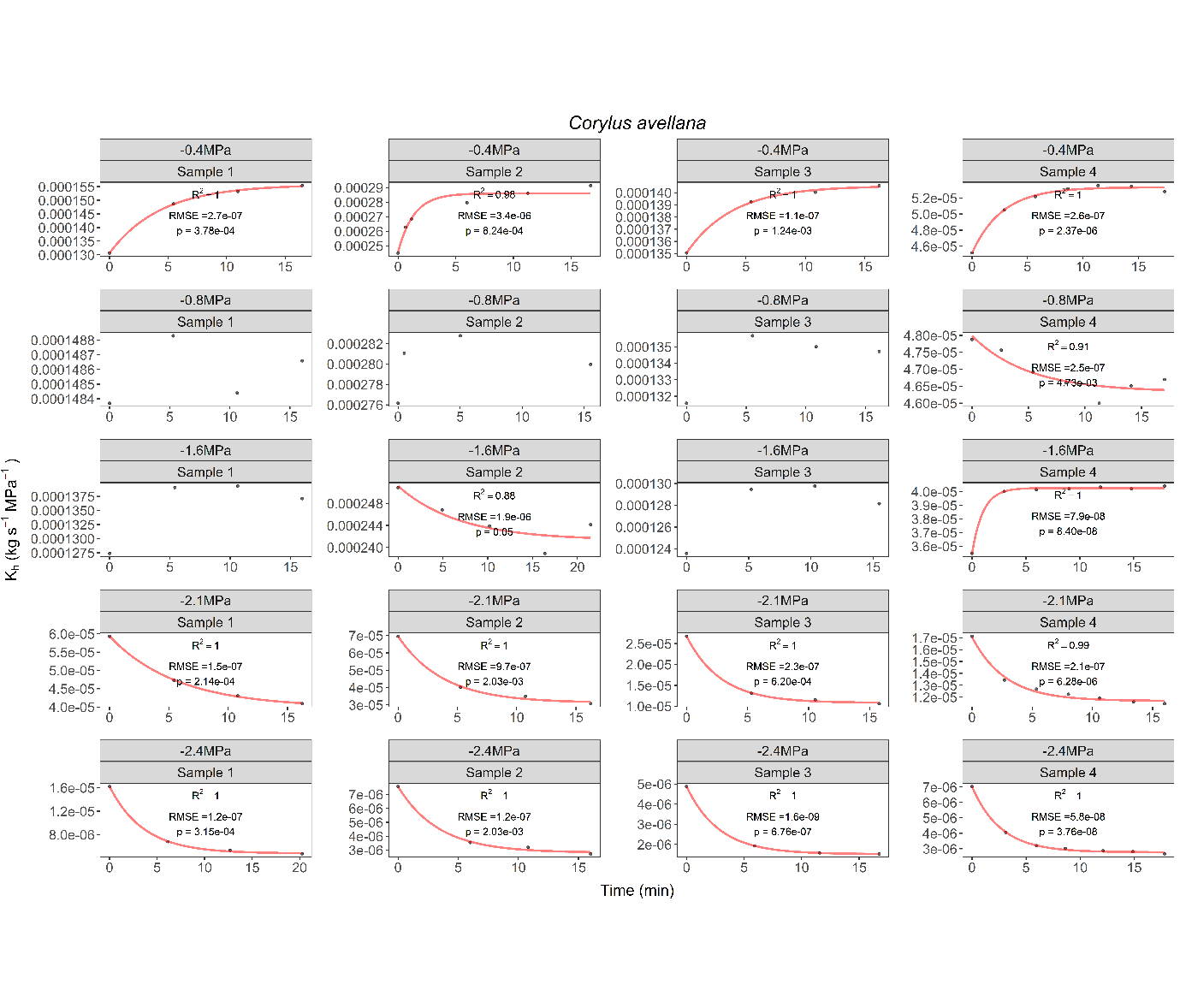

**Fig. S11** Measured (dots) vs. predicted (full red line) hydraulic conductivity (K_h_) values over time for four samples of *Corylus avellana*, five water potentials (Ψ), and temperatures of 22^o^C. Dots represent stem samples that were spun in a flow-centrifuge for 15 minutes at a constant temperature and subjected to a constant Ψ based on a given rotational speed. The curves were predicted values of K_h_, which were estimated by an asymptotic exponential model. The root mean squared error (RMSE), R-squared (R^2^), and p-values (p) were used as a measure of the goodness of fit.

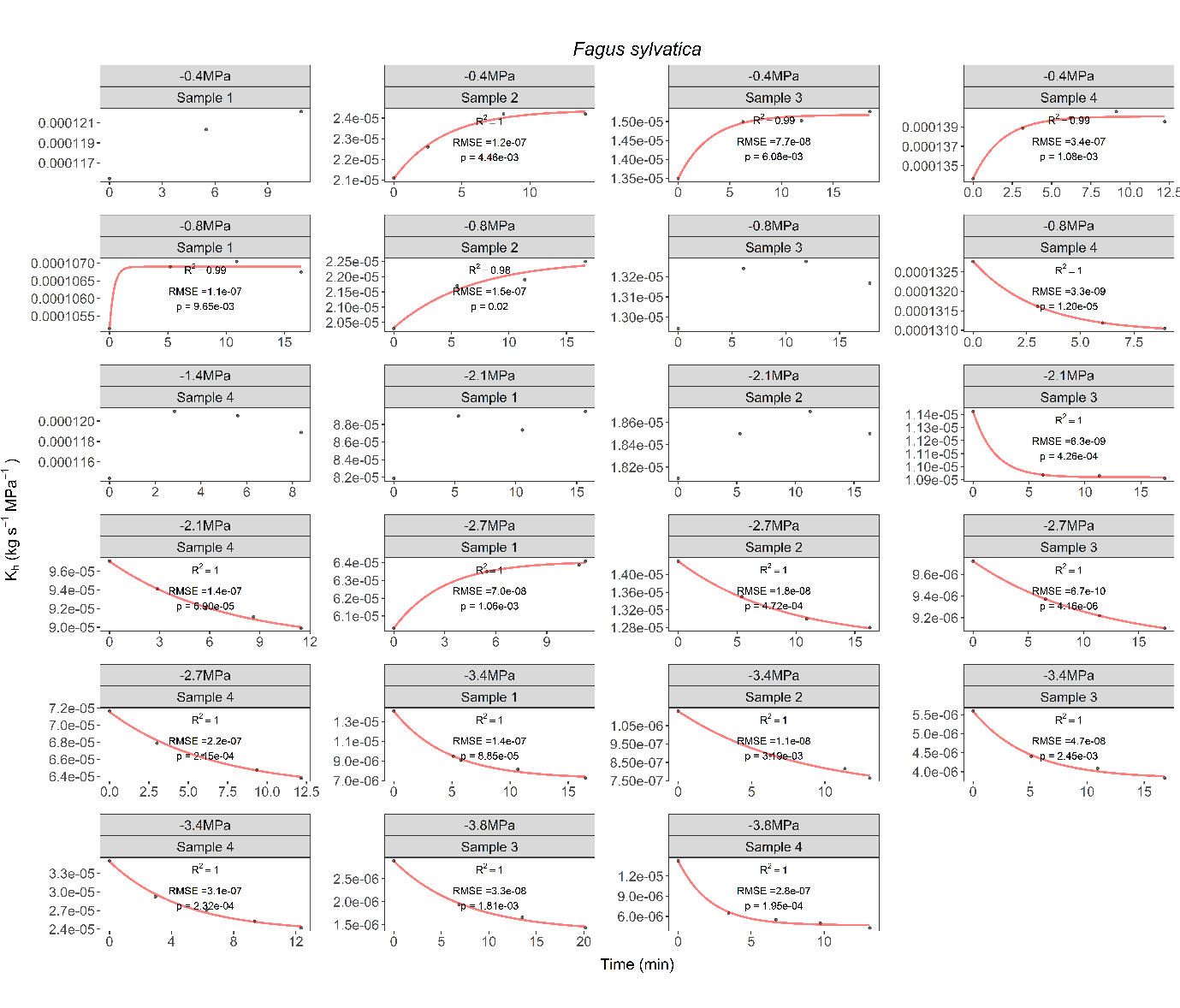

**Fig. S12** Measured (dots) vs. predicted (full red line) hydraulic conductivity (K_h_) values over time for four samples of *Fagus sylvatica*, seven water potentials (Ψ), and temperatures of 22^o^C. Dots represent stem samples that were spun in a flow-centrifuge for 15 minutes at a constant temperature and subjected to a constant Ψ based on a given rotational speed. The curves were predicted values of K_h_, which were estimated by an asymptotic exponential model. The root mean squared error (RMSE), R-squared (R^2^), and p-values (p) were used as a measure of the goodness of fit.

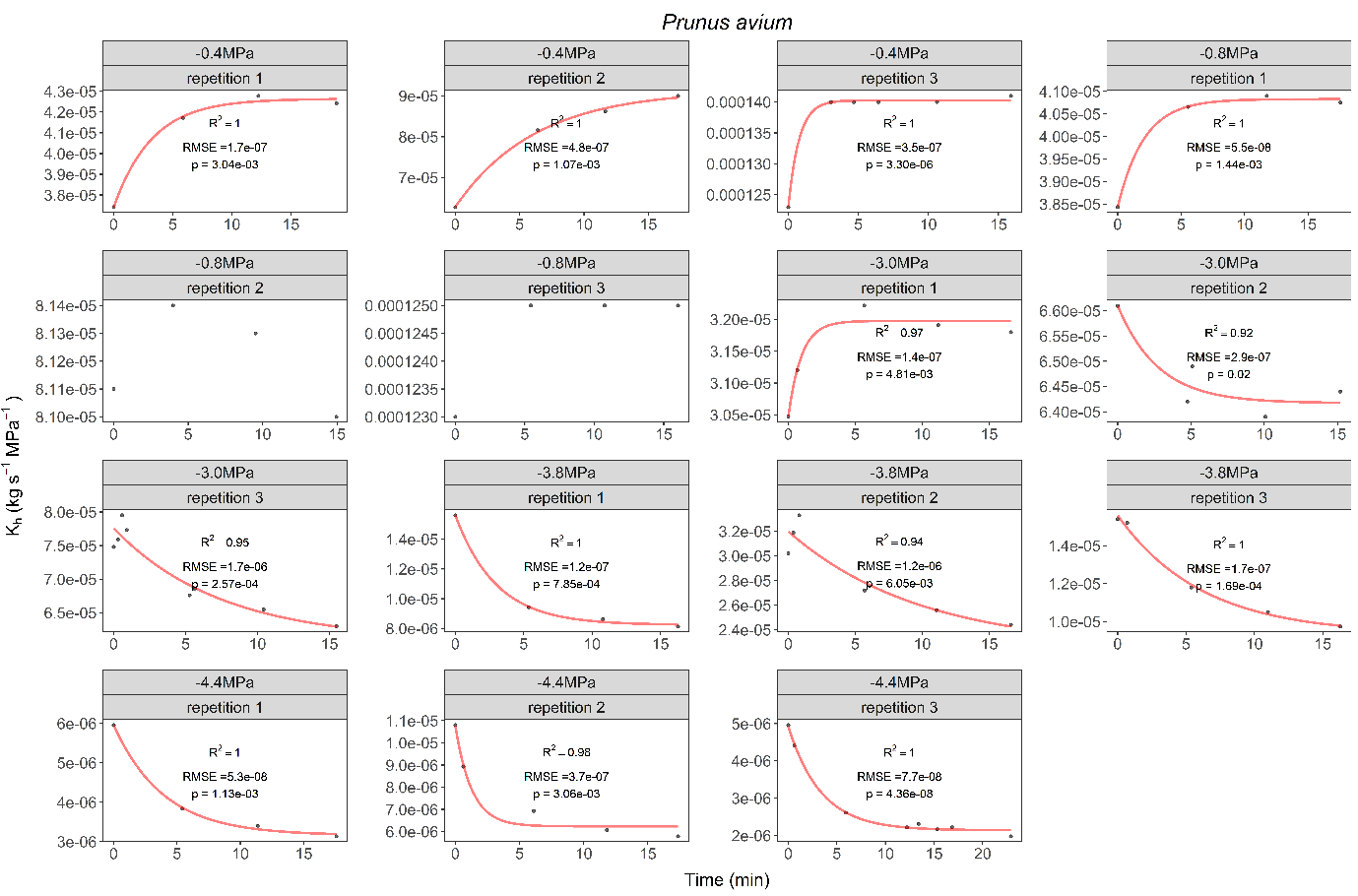

**Fig. S13** Measured (dots) vs. predicted (full red line) hydraulic conductivity (K_h_) values over time for four samples of *Prunus avium*, five water potentials (Ψ), and temperatures of 22^o^C. Dots represent stem samples that were spun in a flow-centrifuge for 15 minutes at a constant temperature and subjected to a constant Ψ based on a given rotational speed. The curves were predicted values of K_h_, which were estimated by an asymptotic exponential model. The root mean squared error (RMSE), R-squared (R^2^), and p-values (p) were used as a measure of the goodness of fit.

**Methods S1:** The application of an asymptotic exponential model for estimating time-stable hydraulic conductivity.

The asymptotic exponential model, as a nonlinear regression, was preferred over polynomial regression due to its applicability to biological and physical phenomena (Bates & Watts, 1988), particularly in scenarios involving complex and nonlinear processes. Notably, these models offer more interpretable parameters, as they can be associated with biologically meaningful processes (Miguez et al., 2018). For instance, the asymptotic parameters (y_asym_ and $a$) and intercept (*c*) had units equal to the response variable (*y*), and the initial slope of the curve (*b*) had units equal to the response variable divided by the independent variable (*x*).

According to Spitters, (1986), the asymptotic exponential equation could be expressed as:

$$y(x)=a(1-e^{\frac{-bx}{a}}) + c$$

(4)

By considering the limit as *x* tended to infinity in equation 4, it was possible to calculate the *y* value in the asymptote (*y_asym_*):

$y_{asym} \underset{x\to\infty}{= lim} [a(1-e^{\frac{-bt}{a}}+c)]=a$ $+ c$

(5)

Utilizing this approach, (Silva et al., 2023) employed this model to describe the decline of K_h_ over time and to estimate the lowest value of hydraulic conductivity (K_low_). To accomplish this, the authors applied a negative transformation to equation 4. Therefore, considering *x* = t, *y* = K_h_, *c* = K_0_, and *y*_asym_ = K_low_ equations 4 and 5 could be expressed as follows:

$$K_{h}(t)=-a(1-e^{\frac{-bt}{a}}) + K_{0}$$

(6)

$$K_{\mathrm{low}}=- a +K_{0}$$

(7)

Here, we utilized equations 6 and 7 to describe the decrease in K_h_ and estimated the time-stable K_h_ in experiments 1 and 2. Similarly, we applied the same methodology to describe the increase in K_h_. In this context, no transformation of equation 4 was necessary. Hence, the expressions for K_h_(t) increasing and the highest hydraulic conductivity values (K_high_) can be formulated as follows:

$$K_{h}(t)=a(1-e^{\frac{-bt}{a}}) + K_{0}$$

(8)

$$K_{\mathrm{high}}=a +K_{0}$$

(9)

To simplify the notation, we adopted K_asym_ as the time-stable K_h_ value, which was in the asymptote of the curve, without distinguishing whether K_asym_ corresponded to K_low_ or K_high_. We developed a script that automatically identified the direction of the K_h_(t) curve. This script estimated parameters such as K_0_, $a$, and *b* (see Fig. S1), and returned the most likely K_h_ value in the asymptote, which could be either K_high_ or K_low_ (which was not critical for our purposes as we only needed a time-stable value). Additionally, it was essential to elucidate the difference between K_max_ and K_0_. K_max_ represented the maximum measured K_h_ value, used to calculate PLC for VCs (equation 1). K_0_ is an estimated value of K_h_ at time zero when assessing the time effect at a fixed Ψ.

All analyses and nonlinear regressions were developed using the R programming environment. Codes are available upon request.

**References**

Bates, D. M., & Watts, D. G. (1988). *Nonlinear regression analysis and its applications*. Wiley.

Guan, X., Werner, J., Cao, K. F., Pereira, L., Kaack, L., McAdam, S. A. M., & Jansen, S. (2022). Stem and leaf xylem of angiosperm trees experiences minimal embolism in temperate forests during two consecutive summers with moderate drought. *Plant Biology*, *24*(7), 1208–1223. https://doi.org/10.1111/plb.13384

Miguez, F., Archontoulis, S., & Dokoohaki, H. (2018). *Chapter 15: Nonlinear Regression Models and Applications* (B. G. Yeater & K. M., Eds.). Applied Statistics in Agricultural, Biological, and Environmental Sciences. https://doi.org/10.2134/appliedstatistics.2016.0003

Silva, L. M., Pereira, L., Kaack, L., Guana, X., Trabia, C. L., & Jansen, S. (2023). Gas diffusion kinetics drive embolism spread in angiosperm xylem: evidence from flow- centrifuge experiments and modelling. *BioRxiv*, doi: 10.1101/2023.04.19.537442.

Spitters, C. J. T. (1986). Separating the diffuse and direct component of global radiation and its implications for modeling canopy photosynthesis part ii. Calculation of canopy photosynthesis. *Agricultural and Forest Meteorology*, *38*(1–3), 231–242.

Wheeler, J. K., Sperry, J. S., Hacke, U. G., & Hoang, N. (2005). Inter-vessel pitting and cavitation in woody Rosaceae and other vessel led plants: A basis for a safety versus efficiency trade-off in xylem transport. *Plant, Cell and Environment*, *28*(6), 800–812. https://doi.org/10.1111/j.1365-3040.2005.01330.x
